# The Drosophila Afadin and ZO-1 homologs Canoe and Polychaetoid act in parallel to maintain epithelial integrity when challenged by adherens junction remodeling

**DOI:** 10.1101/609123

**Authors:** Lathiena A. Manning, Kia Z. Perez-Vale, Kristina N. Schaefer, Mycah T. Sewell, Mark Peifer

## Abstract

During morphogenesis cells must change shape and move without disrupting tissue integrity. This requires cell-cell junctions to allow dynamic remodeling while resisting force generated by the actomyosin cytoskeleton. Multiple proteins play roles in junctional-cytoskeletal linkage, but the mechanisms by which they act remain unclear. Drosophila Canoe maintains adherens junction-cytoskeletal linkage during gastrulation. Canoe’s mammalian homolog Afadin plays similar roles in cultured cells, working in parallel with ZO-1 proteins, particularly at multicellular junctions. We took these insights back into the fly embryo, exploring how cells maintain epithelial integrity when challenged by adherens junction remodeling during germband extension and dorsal closure. We found Canoe helps cells maintain junctional-cytoskeletal linkage when challenged by the junctional remodeling inherent in mitosis, cell intercalation and neuroblast invagination, or by forces generated by the actomyosin cable at the leading edge. However, even in the absence of Canoe many cells retain epithelial integrity. This is explained by a parallel role played by the ZO-1 homolog Polychaetoid. In embryos lacking both Canoe and Polychaetoid, cell junctions fail early, with multicellular junctions especially sensitive, leading to widespread loss of epithelial integrity. Our data suggest Canoe and Polychaetoid stabilize Bazooka/Par3 at cell-cell junctions, helping maintain balanced apical contractility and tissue integrity.

## Introduction

Building the animal body and maintaining tissue homeostasis require the coordinated effort of many cells acting in concert. Cells must change shape and move but need to do so without disrupting tissue integrity. These paired needs require integration of the cell adhesion and actomyosin cytoskeletal machinery, which work together to provide cells, tissues, and organs with the correct architecture and allow them to change shape and move in coordinated ways (Heer and Martin, 2017). Epithelial cells, a polarized cell type that act as the building blocks for most tissues, must coordinate adhesion and the cytoskeleton during tissue development. These cells are organized into sheets with apical-basal polarity and are connected by intercellular adhesion complexes. Cadherin-based adherens junctions (AJs) provide connections between cells and form the boundary between the apical and basolateral domains. Transmembrane cadherins mediate cell-cell adhesion, while p120-catenin, β-catenin and α-catenin, bound to the cadherin cytoplasmic tails, stabilize cadherins at the cell surface and interact with the actomyosin cytoskeleton (Meng and Takeichi, 2009; Mege and Ishiyama, 2017). Disruption or dysregulation of AJs leads to disorganization of tissue architecture, which is a common step in solid tumor metastasis and numerous developmental disorders.

These vital roles of AJs have made them the subject of intensive research. In the conventional model, cadherins linked directly to actin via α- and β-catenin (Rimm *et al*., 1995; Ozawa, 1998). However, more recent work revealed that this linkage is mediated by a far more sophisticated set of molecules (Mege and Ishiyama, 2017). This led to the search for additional linker proteins that regulate epithelial cell adhesion and AJ/cytoskeletal linkage. One such junction-linker protein is fly Canoe (Cno) and its mammalian homolog Afadin. Cno’s multidomain structure allows it to interact directly with the cytoskeleton via its F-actin-binding domain and to bind AJ proteins, including E-cadherin and α-catenin, via its PDZ and proline-rich domains (Miyamoto *et al*., 1995; Mandai *et al*., 2013).

We initially hypothesized that *Drosophila* Cno would be essential for cell adhesion, as was observed for E-cadherin (Ecad) (Tepass *et al*., 1996), α– (Sarpal *et al*., 2012) and β-catenin (Cox *et al*., 1996). However, to our surprise, *cno* maternal/zygotic mutants (*cnoMZ*) maintain epithelial integrity throughout gastrulation (Sawyer *et al*., 2009), unlike embryos lacking Ecad or the catenins. Instead, our analysis revealed that while Cno is not essential for maintaining cell-cell adhesion, it is required for many morphogenetic movements requiring AJ/cytoskeletal linkage, including apical constriction and subsequent internalization of the mesoderm, effective cell intercalation during germband elongation, and dorsal closure (Sawyer *et al*., 2009; Choi *et al*., 2011; Sawyer *et al*., 2011; Miyamoto *et al*., 1995; Boettner *et al*., 2003). In the absence of Cno, the actomyosin cytoskeleton detaches from AJs, consistent with a role as a linker. This is particularly striking during germband extension, which is largely driven by coordinated, opposing planar polarity of AJs/Bazooka (Baz; the fly Par3 homolog) and the actomyosin cytoskeleton, promoting polarized contractility across the entire tissue in the direction of elongation (reviewed in Vichas and Zallen, 2011; Harris, 2018). Loss of Cno enhances AJ and Baz planar polarity to dorsal-ventral cell boundaries and simultaneously leads to retraction of the actomyosin cytoskeleton from the anterior-posterior cortex (Sawyer *et al*., 2011). *cno* and *baz* mutants exhibit strong genetic interactions, consistent with a mechanistic connection (Sawyer *et al*., 2011). An additional role for Cno in later epidermal integrity is suggested by its cuticle phenotype, with the ventral epidermis most sensitive to disruption (Sawyer *et al*., 2009). This special sensitivity of the ventral epidermis is shared in several situations involving reduction in AJ (Tepass *et al*., 1996) or apical polarity proteins (Harris and Tepass, 2008). We explore the mechanistic basis for Cno’s role in this process here.

Another potential set of junction-cytoskeletal linker proteins are those of the Zonula occludens (ZO-1) family. Like Cno, these are multidomain scaffolding proteins that can directly bind F-actin and bind a wide variety of junctional proteins (Fanning and Anderson, 2009). ZO-1 family members are best known for their roles in tight junctions, which in mammals localize just apical to the AJ (Van Itallie and Anderson, 2014). In tight junctions, strands of claudins are positioned apically and cross-linked to the actin cytoskeleton by ZO-1, providing epithelial barrier function. In the absence of ZO-1 family function, claudin strands disperse all along the lateral membrane and barrier function is disrupted (Umeda *et al*., 2006). ZO-1 family proteins also localize to nascent AJs where they are thought to have roles in accelerating AJ assembly (Ikenouchi *et al*., 2007; Yamazaki *et al*., 2008). Mice have three ZO family members, with partially overlapping expression patterns; loss of function mutants in ZO-1 (Katsuno *et al*., 2008) and ZO-2 (Xu *et al*., 2008) have distinct embryonic lethal phenotypes, suggesting partial redundancy. *Drosophila* has only a single family member, Polychaetoid (Pyd; (Takahisa *et al*., 1996). However, *Drosophila* lacks apical tight junctions, and Pyd localizes to AJs throughout development (Wei and Ellis, 2001; Jung *et al*., 2006; Seppa *et al*., 2008; Choi *et al*., 2011). We were surprised to learn that *pyd* maternal/zygotic null mutants can survive to adulthood, with defects in Notch signaling that affect bristle development (Choi *et al*., 2011; Djiane *et al*., 2011). However, 60% of maternal/zygotic mutant embryos die, with defects in cell shape change during dorsal closure and defects in tracheal development. Thus, neither Cno or Pyd alone are essential for early epithelial integrity. Intriguingly, although Afadin and ZO-1 localize to distinct though adjacent junctions in mammalian cells, they can physically interact with each other in both mammals and *Drosophila* and early studies of weak alleles in *Drosophila* indicate a potential synergistic interaction (Yamamoto *et al*., 1997; Takahashi *et al*., 1998). However, since neither allele used in these experiments was a null allele, it was impossible to distinguish whether Cno and Pyd work together in the same process or work in parallel.

Our studies of Cno’s homolog Afadin in mammalian MDCK cells provided another set of insights (Choi *et al*., 2016). In these cells, reducing Afadin levels has only subtle effects. Reducing levels of ZO-1 family members, in contrast, stimulates robust assembly of a contractile actomyosin array at the apical adherens junction (Fanning *et al*., 2012), via activation of Shroom and Rho kinase (Choi *et al*., 2016). In these cells, each cell border, bounded by tricellular junctions, serves as an independent contractile unit. Borders are contractile, but within homeostatic limits, as balanced contractility between different cell borders maintains individual cell borders at roughly similar lengths and thus cell shape is relatively homogenous. Knockdown of Afadin in this ZO-1 knockdown background strikingly disrupted this homeostatic balance, leading to highly unbalanced contractility. Some cell borders became shortened and hypercontractile while others became hyper-elongated. Disruptions in the actomyosin cytoskeleton at cell junctions were most readily apparent at tricellular and multicellular junctions, where the tight bundling of actin and myosin in the AJs was lost. However, these disruptions could spread into neighboring bicellular borders. These data suggested the possibility that Afadin and Cno may play an additional role in helping maintain balanced contractility on different cell borders, and thus maintain epithelial integrity.

Cno’s diverse functions in embryonic development mean that the early effects of its loss make it challenging to assess whether effects later are primary or secondary. We thus developed RNAi tools to reduce Cno function to different extents. We also developed methods to simultaneously reduce the function of Cno and Pyd, to explore whether they act together or in parallel in epithelial tissues. This revealed important roles for Cno in balancing contractility on different cell borders throughout development and suggests that Cno and Pyd/ZO-1 act in parallel in maintaining adhesion and junctional integrity during morphogenesis.

## RESULTS

### Developing tools to titrate reduction of Canoe function, allowing exploration of its full range of roles in embryonic development

*cno* was originally identified in *Drosophila* through the effect of zygotic mutants on dorsal closure (Jürgens *et al*., 1984; Miyamoto *et al*., 1995; Choi *et al*., 2011). The zygotic null mutant has relatively mild defects during this process, since these embryos retain at least some maternally-contributed Cno through dorsal closure (Choi *et al*., 2011). Later studies of maternal/zygotic loss of *cno* (*cnoMZ*), in which Cno expression is completely removed, revealed important roles of Cno in mesoderm invagination and germband elongation (Sawyer *et al*., 2009; Sawyer *et al*., 2011). These studies also suggested that Cno regulates the link between AJ and actin during apical constriction. We suspected Cno also played important subsequent roles. Analysis of the *cnoMZ* cuticle phenotype (Sawyer *et al*., 2009) suggested a later role in epidermal integrity, but the mechanisms by which it acts in maintaining epidermal integrity were not known. In addition, the severity of the *cnoMZ* terminal phenotype made studying its role in late embryonic events like dorsal closure difficult, as it is hard to distinguish between primary and secondary consequences of Cno loss.

To explore Cno’s roles in the full set of developmental events in which it is involved, we hypothesized that utilizing RNA interference (RNAi) in conjunction with the Gal4-UAS system (Brand and Perrimon, 1993; Duffy, 2002) would allow us to titrate Cno knockdown to different levels in order to study a wider variety of post-gastrulation events. The TRiP project has generated lines expressing shRNAs under the control of Gal4 drivers against many *Drosophila* genes (Perkins *et al*., 2015), including *cno* (Bonello *et al*., 2018), allowing efficient knockdown. The community has also generated a wide variety of lines expressing GAL4 in different tissues and at different times. Among these are lines expressed during mid-oogenesis, allowing knockdown of maternally-contributed mRNAs, and continued knockdown of zygotic mRNAs in the progeny (Staller *et al*., 2013). The strongest of these can result in a maternal/zygotic null phenotype. In an effort to obtain different degrees of Cno knockdown we generated females carrying one of three different maternal Gal4 drivers along with the *UAScnoRNAiValium20shRNA* or *UAScnoRNAiValium22shRNA* constructs, and tested their phenotypes. Our tests ordered these maternal drivers into the relatively weak *nos-Gal4*, the moderate *MTD-Gal4 driver*, and the strong “*mat-Gal4-2+3*” driver, carrying a maternal α-tubulin-GAL4 driver on both the second and third chromosomes (*shRNA* and *GAL4* lines used are described in detail in Table 2)

As an initial screen of how different degrees of Cno knockdown affect morphogenesis, we assessed embryo lethality and cuticle phenotype, as the latter reveals the success of major morphogenetic movements and the effect on epidermal integrity. We created categories to illustrate the range of morphogenic phenotypes seen in different *cno* mutant or knockdown genotypes (Fig. 1A-I). Head involution is most sensitive to Cno reduction (Fig. 1A-C), with defects in dorsal closure seen after moderate reduction (Fig. 1D-F), and finally defects in epidermal integrity observed in the strongest mutant combinations (Fig. 1G-I). As our baselines for comparison, we used *cno* zygotic null mutants (zygotic *cno^R2^*/*cno^R2^*; Sawyer *et al*., 2009), which retain maternally contributed Cno, and maternal/zygotic *cno^R2^* null mutants (*cnoMZ;* Sawyer *et al*., 2009), which completely lack Cno. Zygotic null mutants exhibit fully penetrant embryonic lethality, but defects in morphogenesis are relatively mild, ranging from mild to strong defects in head involution (Fig. 1K; Table 1; Sawyer *et al*., 2009). At the other end of the phenotypic spectrum, *cnoMZ* mutants also exhibit complete embryonic lethality (Fig. 1J; n=432) but cuticle defects are much more severe, with most embryos exhibiting complete failure of both head involution and dorsal closure and many with more severely disrupted epidermal integrity (Fig. 1K; Table 1; Sawyer *et al*., 2009).

**Figure 1.**
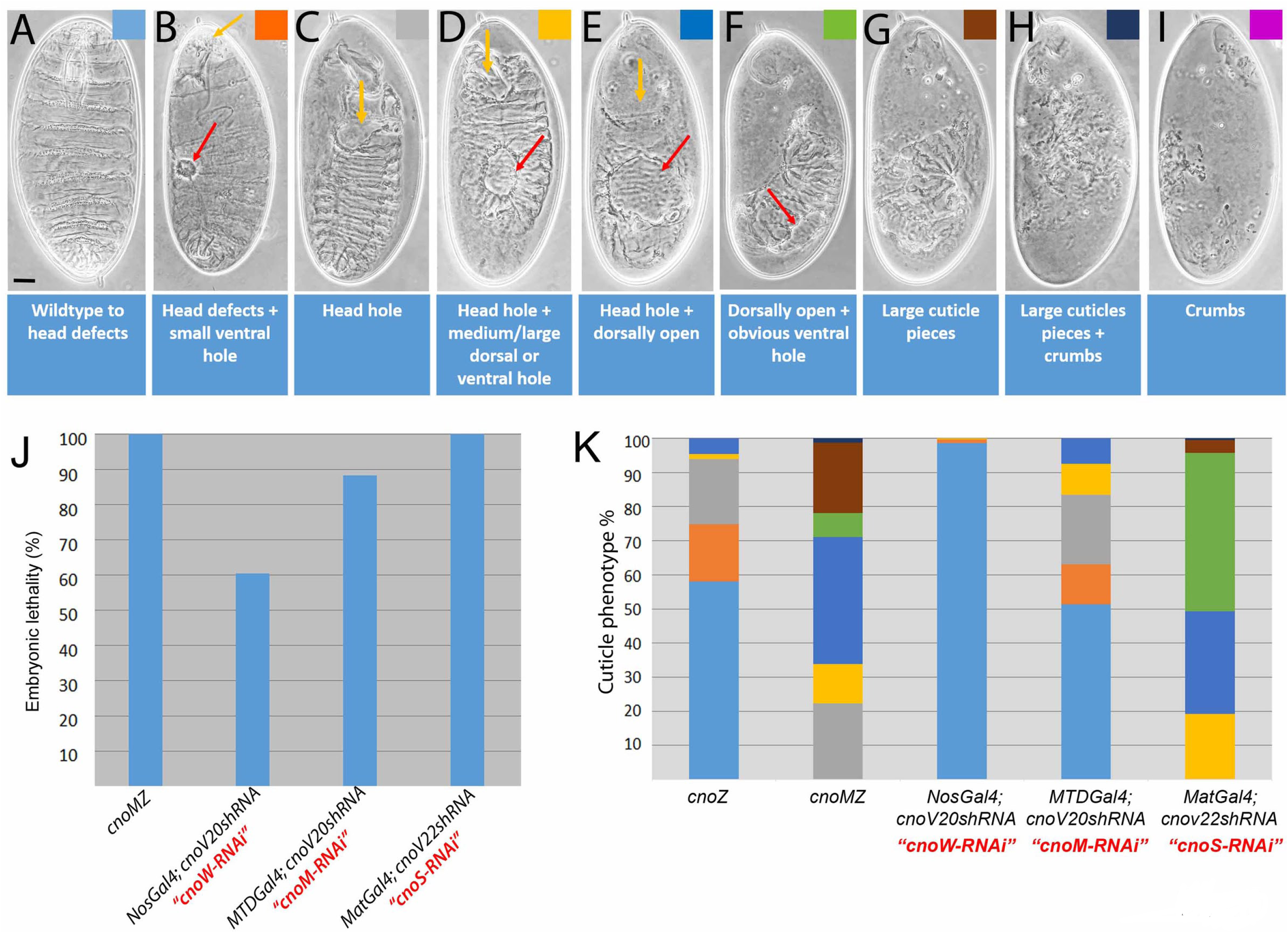
Developing tools to modulate Cno function using RNAi. A-I. Cuticle preparations revealing the spectrum of defects in morphogenesis and epithelial integrity seen in embryos with different degrees of Cno function. A. Nearly wildtype cuticle with subtle defects in the head skeleton (arrow). Scale bar = 30µm. B. Defects in head involution are accompanied by small holes in the ventral or dorsal cuticle (arrow). C. Failure of head involution, leaving a hole in the anterior (arrow). D. Failure of head involution (yellow arrow) accompanied by a large hole in the ventral or dorsal cuticle (red arrow). E. Failure of both head involution (yellow arrow) and dorsal closure (red arrow). F. Dorsally open with hole in the remaining cuticle (arrow). G. Remaining cuticle in large fragments. H. Large and small fragments of cuticle (crumbs-like; yellow arrow). I. Only small cuticle fragments remaining. J. Embryonic lethality of different *cno* RNAi genotypes relative to *cno^R2^* maternal/zygotic mutants. K. Severity of cuticle phenotype of different *cno* RNAi genotypes relative to *cno^R2^* zygotic or maternal/zygotic mutants.

**Table 1:** Cuticle Phenotypes

| Genotype | wildtype or minor head defects | Head defects + small ventral hole | Head hole | Head hole + medium/ large dorsal or ventral hole | Head hole + dorsally open | totally dorsal open + obvious ventral hole | Large cuticle pieces | Large cuticle pieces + crumbs | Crumbs only | total | Lethality % |
| --- | --- | --- | --- | --- | --- | --- | --- | --- | --- | --- | --- |
| *cno<sup>R2</sup>* zygotic null (*CnoZ*) | 58% | 17% | 19% | 1.50% | 5.00% | 0 | 0 | 0 | 0 | 131 | 100% |
| *cno<sup>R2</sup>* maternal and zygotic (MZ) null | 3% | 1% | 21% | 11% | 36% | 7% | 20% | 1% | 0 | 253 | 100% |
| NosGal4; *cnoV20shRNA* (weak = *cnoW-RNAi*) | 99% | 1% | 0 | 0.30% | 0 | 0 | 0 | 0 | 0 | 367 | 61% |
| MTDGal4; *cnoV20shRNA* (moderate = *cnoM-RNAi*) | 51% | 12% | 20% | 9% | 7.50% | 0 | 0 | 0 | 0 | 333 | 88% |
| MatGal4; *cnoV20shRNA* (control cross for double mutant) | 21% | 2% | 18% | 23% | 25% | 8% | 2% | 2% | 0 | 665 | 100% |
| MatGal4; MatGal4/*cnoV22shRNA* (strong = *cnoS-RNAi*) | 0 | 0 | 0.20% | 19% | 30% | 46% | 4% | 0.40% | 0 | 521 | 84% |
| *pyd<sup>B12</sup>/pyd<sup>B12</sup>* *cnoV20shRNA* (*pyd cno* mutant) | 0.20% | 0 | 5% | 13% | 13% | 22% | 23% | 19% | 5% | 471 | 100% |

Our different *cno* RNAi strategies delivered a wide range of phenotypes, from embryos with no observable defects to those with severe morphogenic defects like those seen in *cnoMZ*. The vast majority of embryos in which our weak *nos-Gal4* drove *UASCnoRNAi*2*0shRNA* (*cnoW-RNAi)* exhibited largely normal cuticle morphology, with virtually all having only minor head defects (Fig. 1K; Table 1), and 39% of the embryos are viable (n=171; Fig. 1J). The moderate *MTDGal4* driving *UASCnoRNAi*20*shRNA* (referred to below as *cnoM-RNAi*) led to more penetrant embryonic lethality (88%, n=330; Fig. 1J), and provided the broadest spectrum of cuticle phenotypes. Half of the progeny had mild head defects and half had strong disruption of head involution and mild effects on dorsal closure (Fig. 1K; Table 1). The strongest GAL4/RNAi combination, *mat-Gal4-2+3* driving *UASCnoRNAi*2*2shRNA* (below referred to as *cnoS-RNAi*), had completely penetrant lethality (100%; n=389; Fig. 1J) and the strongest embryonic defects, with 30% exhibiting the “canoe” phenotype, reflecting complete failure of head involution and dorsal closure (Fig. 1F), and 50% having additional defects in epidermal integrity (Fig. 1K). *cnoS-RNAi* embryos had Cno reduced to essentially undetectable levels at the onset of development (to 1.3% of wildtype; Suppl. Fig. 1A,B), and Cno levels remained drastically reduced at the end of morphogenesis (to 4.9% of wildtype; Suppl. Fig. 1A,C; Analysis by immunofluorescence confirmed these reductions; Suppl. Fig. 1G’ vs. H’). *cnoS-RNAi* largely phenocopied the maternal/zygotic loss of Cno, as assessed by cuticle pattern (Fig. 1K). Together, this set of UAS-RNAi/GAL4 lines provided us with the ability to dial Cno function down to the desired degree to study different embryonic events. In subsequent analysis we used *cnoS-RNAi* to analyze earlier embryonic events beginning during and just after germband extension, allowing us to explore the genesis of epithelial integrity defects, and focused on the moderate *cnoM-RNAi* in studies examining the role of Cno during dorsal closure.

### Canoe is required to maintain homeostatic cell shapes and balanced contractility along the leading edge during dorsal closure

Our recent super-resolution imaging in cultured mammalian MDCK cells suggested Cno’s homolog Afadin plays a prominent role at tricellular junctions where three cells meet, reinforcing end-on links between the actomyosin cytoskeleton and cell-cell junctions and thus allowing tricellular junctions to resist mechanical tension generated by actomyosin contractility along the cell border (Choi *et al*., 2016). These data fit with our earlier work in *Drosophila*, which revealed Cno is enriched at tricellular junctions and strengthens actomyosin-junction linkages during mesoderm invagination and germband elongation (Sawyer *et al*., 2009; Sawyer *et al*., 2011; Bonello *et al*., 2018). We sought to further explore Cno’s role at tricellular junctions in in vivo. Dorsal closure provides a superb place to study this, as the dorsalmost cells of the lateral epidermis assemble a planar-polarized contractile actomyosin cable at their leading edge, anchored cell-to-cell at tricellular junctions. Contraction of this supercellular cable along with pulsed contractions of the more dorsal amnioserosal cells help power closure (reviewed in Hayes and Solon, 2017; Kiehart *et al*., 2017). Previous analyses of Cno’s role in dorsal closure relied on zygotic *cno* mutants (e.g. Choi *et al*., 2011; Boettner *et al*., 2003). However, most zygotic null mutants complete closure due to maternally contributed protein (Sawyer *et al*., 2009; Fig. 1), and thus the phenotype we observed does not reveal Cno’s full role during this process.

Our new set of RNAi tools allowed us to dial down Cno function to a level at which dorsal closure could be initiated, but in which Cno function was substantially lower than that seen in zygotic *cno* mutants (Fig. 1K). To do so, we drove the *cnoV20shRNA* with the Maternal Triple Driver (MTD)-GAL4 (Mazzalupo and Cooley, 2006); *cnoM-RNAi)*. Embryos that received two copies of the hairpin had substantial defects in head involution and dorsal closure (Fig. 1A-K). We thus began by examining embryos during these stages stained with antibodies against AJ and cytoskeletal proteins, to allow us to assess morphogenetic movements and cell shape change during dorsal closure.

Wildtype dorsal closure begins at the completion of germband retraction. The scalloped boundary between the epidermis and amnioserosa (Fig. 2A, red arrows) straightens (Fig. 2B,I, red arrows) as the actin cable is assembled. Epidermal cells elongate along the dorsal-ventral axis, beginning with those at the leading edge (Fig. 2A, inset) and then the cells more ventrally (Fig. 2B,I). The forces generated by amnioserosal cell apical constriction and the contractile actomyosin cable assembled at the leading edge (LE) combine to gradually close the dorsal opening (Kiehart *et al*., 2000; Hutson *et al*., 2003); Fig. 2B-D). As the two sheets meet at the anterior and posterior canthi, they zipper together (Fig. 2C-D, red arrows) until the opening is closed and the embryo is completely enclosed in epidermis. At that point the amnioserosal cells undergo apoptosis (Toyama *et al*., 2008). Head involution occurs in parallel (Fig 2B-D, yellow arrows). In contrast, morphogenesis was severely altered in *cnoM-RNAi* embryos. They exhibited strong defects in head involution, with the head epidermis lost or disrupted (Fig. 2E-H vs B-D, yellow arrows), and germband retraction was not completed (Fig2F,G vs B,C, green arrows). Some defects observed were similar to those previously observed in *cno* zygotic null mutants (Choi *et al*., 2011), like the retention of excessively deep segmental grooves (Fig. 2F-H vs D, blue arrows) and a LE that was wavy rather than straight (Fig. 2J vs. I). However, other defects were much more severe. Unlike wildtype embryos, where zippering of the two epidermal sheets is complete before the amnioserosa undergoes apoptosis, in many *cnoM-RNAi* embryos, the two epidermal sheets remained far apart when the amnioserosa began apoptosis, leaving the underlying muscle and gut tissues exposed (Fig. 2F-H, white arrows).

**Figure 2.**
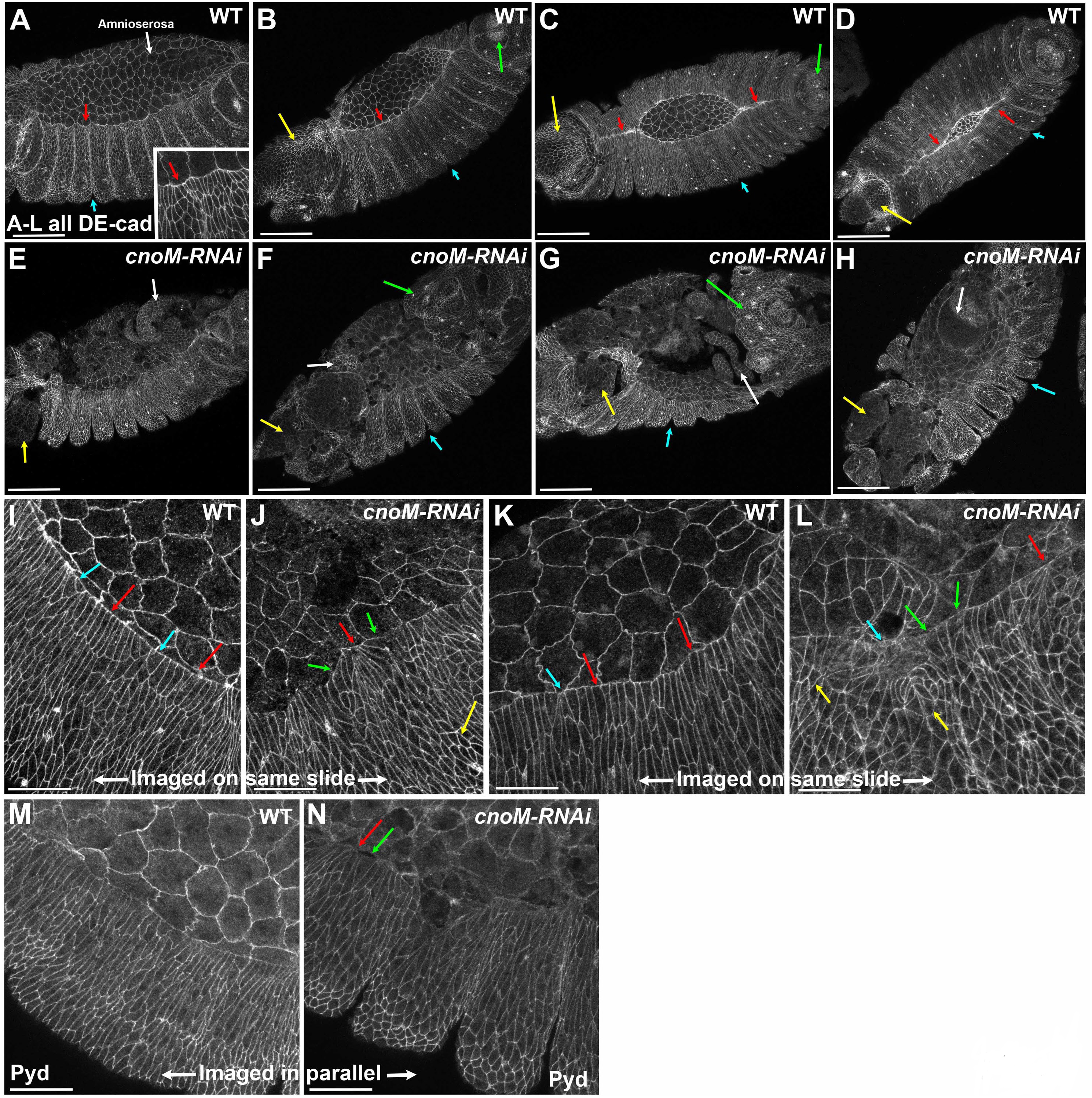
Moderate Cno knockdown disrupts dorsal closure and leads to uneven cell shapes along the leading edge. Embryos, stage 13-14, genotypes indicated. All images show Ecad except M,N which show Pyd. A-D. Wildtype, progression through dorsal closure. A. At onset the leading edge is scalloped (arrows), and segmental grooves remain relatively deep (blue arrow), extending to the leading edge. B-D As closure proceeds the leading edge straightens (B, red arrow). This occurs simultaneously with head involution (yellow arrows). The two epidermal sheets zipper together at the canthi (C,D red arrows). The posterior spiracles have fully retracted to the posterior end (green arrows), and the segmental grooves have largely retracted (blue arrows). E-H. *cnoM-RNAi* embryos at comparable stages. Head involution has failed (yellow arrows). Segmental grooves remain abnormally deep (blue arrows). Germband retraction has not been completed in many embryos (green arrows), and holes appear in the amnioserosa, exposing underlying tissue (white arrows). I-N. Cell shapes at the leading edge. I,K,M. In wildtype cells are largely uniformly elongated along the dorsal-ventral axis and in width along the leading edge (red arrows), with the exception of cells making the segmental grooves (blue arrows). J,L,N *cnoM-RNAi*. Leading edge cell shapes are highly irregular: some cells have hyper-constricted (red arrows) or hyper-elongated (green arrows) leading edges. A subset of cells behind the leading edge fail to elongate along the dorsal-ventral axis (yellow arrows). I vs. J, K vs. L, and M vs. N were imaged on the same slide or in parallel, revealing no dramatic changes in Ecad or Pyd cortical localization. Scale bars: A-H=50 µm and I-N=20µm.

The LE provides a superb place to assess Cno’s mechanisms of action. LE cells assemble a planar polarized actomyosin cable at their dorsal margin, where they contact the amnioserosal cells (Fig. 3A-E). In wildtype the cable initially assembles at the onset of dorsal closure (Fig. 3A) and is maintained and enhanced as closure proceeds (Fig. 3B,C). The cable behaves in a supercellular fashion, exerting force all along its length (Kiehart *et al*., 2000; Hutson *et al.*, 2003). To accomplish this, individual cell cables (Fig. 3D, white arrows) must be connected cell to cell, presumably at cadherin-based AJs (Fig. 3D, blue arrows). Consistent with this, Ecad is particularly enriched at the LE tricellular junctions (Fig. 3E; Kaltschmidt *et al*., 2002), at the location individual cell actin cables are presumably anchored. Cno is also somewhat enriched at these locations (Fig. 3F (Kaltschmidt *et al*., 2002), reminiscent of its enrichment at tricellular junctions earlier in embryogenesis (Sawyer *et al*., 2009; Bonello *et al*., 2018). In our earlier work we also identified a distinctive localization for the actin polymerization regulator Enabled (Ena) at these special tricellular junctions (Gates *et al*., 2007; Choi *et al*., 2011; Nowotarski *et al*., 2014). In vitro, Ena localizes to growing (barbed) ends of actin filaments (reviewed in Edwards *et al*., 2014) Ena is required for effective cell shape change during dorsal closure (Gates *et al*., 2007). Ena is enriched at all tricellular junctions during the extended germband stage (Gates *et al*., 2007). As dorsal closure begins it remains at tricellular junctions of all epidermal cells (e.g., Fig. 4A, yellow arrow), but becomes particularly enriched at LE tricellular junctions (Fig. 4A, blue arrows), and at the borders of segmental groove cells (Fig. 4A, red arrow). As closure proceeds Ena enrichment at the LE tricellular junctions continues to increase (Fig. 4B-C, arrows). The LE thus provides a place to test our hypotheses, based on our earlier work in *Drosophila* embryos and in knockdown MDCK cells, that Cno and Afadin are cytoskeletal-junction crosslinkers that reinforce connections under tension and which may also help balance tension between different borders.

**Figure 3.**
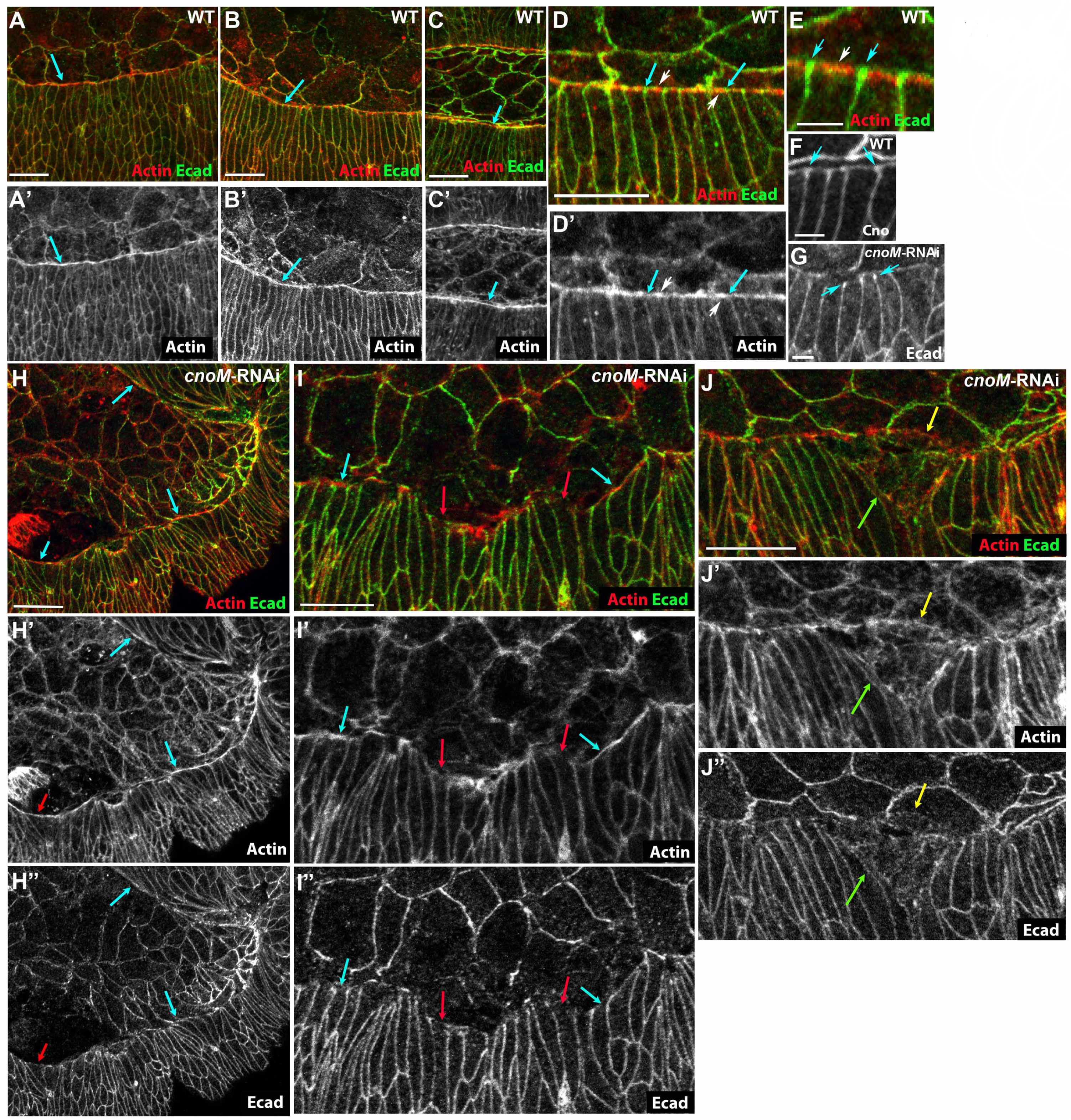
Moderate Cno knockdown does not prevent actin cable assembly but does lead to irregularity in cable maintenance. Embryos, stage 13-14, genotypes indicated. All images show Ecad and actin (imaged with phalloidin) except F which shows Cno. A-F. Wildtype. A-C. Progression through dorsal closure. Actin cable intensifies (blue arrows). D-F. Actin cable (white arrows) is linked cell to cell at leading edge tricellular junctions, where both Ecad and Cno are enriched (blue arrows). G-J. *cnoM-RNAi* embryos. G. Ecad enrichment at leading edge AJs is not lost. H-J The actin cable can be assembled and maintained, even by cells in which the leading edge is hyperextended (blue arrows), but discontinuities in the cable (red arrows) and leading edge cell ripping (J, yellow vs green arrows) were observed. Scale bars in E-G=2µm. All other panel scale bar=10 µm.

**Figure 4.**
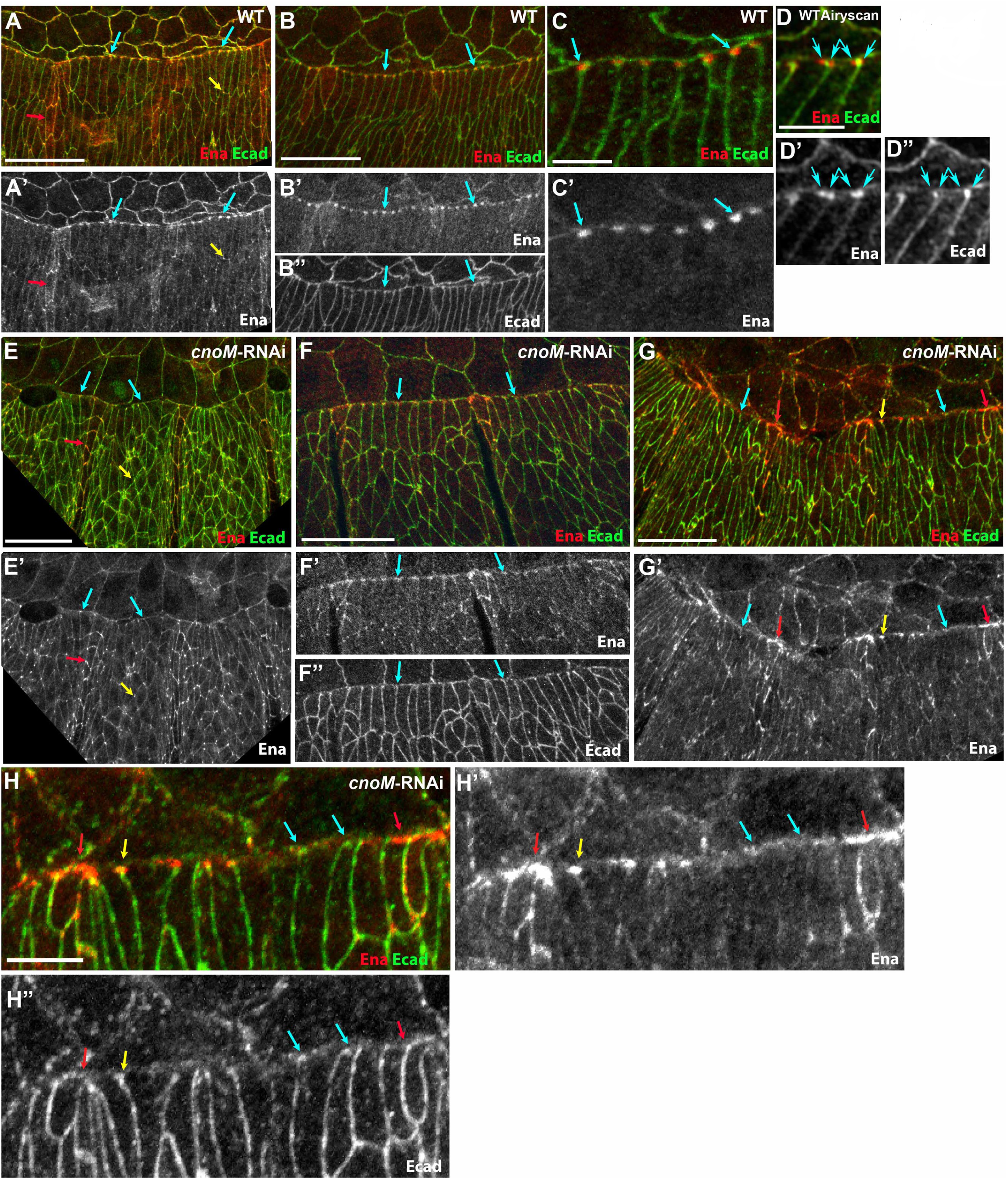
Ena localizes adjacent to leading edge AJs and moderate Cno knockdown disrupts this uniform localization. Embryos, stage 13-14, genotypes indicated. All images show Ecad and Ena. A-D. Wildtype. A. At the onset of dorsal closure Ena localizes to all epidermal tricellular junctions (e.g. yellow arrow), and is especially enriched is segmental groove cells (red arrow). Enrichment near the leading edge AJs begins (blue arrows). B,C. As closure proceeds, enrichment next to leading edge AJs increases. D. Airyscan super-resolution image. Leading edge “dots” sometimes resolve into two dots on the two sides of the Ecad. E-H. *cnoM-RNAi*. E. Early in closure Ena localization to epidermal tricellular junctions (e.g. yellow arrow), segmental groove cells (red arrow) are relatively unchanged from wildtype, while localization near the leading edge AJs is less uniform (blue arrows). F. As closure proceeds, leading edge Ena localization is less focused at tricellular junctions than in wildtype (arrows, compare to B), even when cell shapes are relatively normal. G,H. In most embryos leading edge Ena localization is becomes much more irregular. Distinct leading edge dots remain in some cells (yellow), but at many tricellular junctions Ena localization at leading edge dots is diminished (blue arrows) or broadened (red arrows). Scale bars in A,B,E,F,G=20 µm. Scale bars in C,D, H= 5µm.

We first sought a more detailed understanding of the molecular architecture at the LE, to help us better interpret both the process of wildtype dorsal closure and the effects of reducing Cno. We thus turned to structured illumination microscopy (SIM), using the improved resolution to more precisely define how Cno, Ecad, actin, myosin and Ena are arrayed along the LE relative to one another in 3-dimensions (Fig. 5; our methodology for processing images obtained by SIM is in Suppl. Fig. 2). Standard confocal imaging usually visualized a single Ena “dot” at each tricellular junction (Fig. 4C’), but use of the Zeiss Airyscan module occasionally resolved these into a pair of dots flanking the cadherin-based AJs (Fig. 4D). SIM imaging allowed us to better resolve the Ena “dots”. SIM usually resolved the single Ena dot seen in confocal microscopy into a more complex bipartite structure (Fig. 5B-D), with Ena dots (blue arrows) flanking each side of the Ecad concentrations at the LE tricellular junctions. Viewing these in the Z-axis revealed that Ena precisely aligns with the AJs proteins along the apical-basal axis (Fig. 5E). Imaging Ena together with Cno (Fig. 5F-I) revealed a similar picture, with Ena flanking Cno at LE tricellular junctions (Fig 5G-I, blue arrows). We next imaged Ena along with F-actin (Fig. 5J-M). This revealed that the paired Ena dots (Fig. 5K-L, blue arrows) localized to the tricellular junction-proximal ends of the actin cables (magenta arrows) that run along the LE. Once again, Ena also aligned with the ends of the actin cable along the apical-basal axis (Fig. 5M). Given the known properties of Ena/VASP proteins, this may suggest that actin barbed ends concentrate where the cable interfaces with the cadherin-catenin complex. Finally, our SIM imaging confirmed something that was previously suggested from confocal imaging (e.g., Franke *et al*., 2005): myosin is strongly enriched on the central regions of each actin cable and absent or at reduced levels closest to the LE tricellular junctions (Fig. 5N-R).

**Figure 5.**
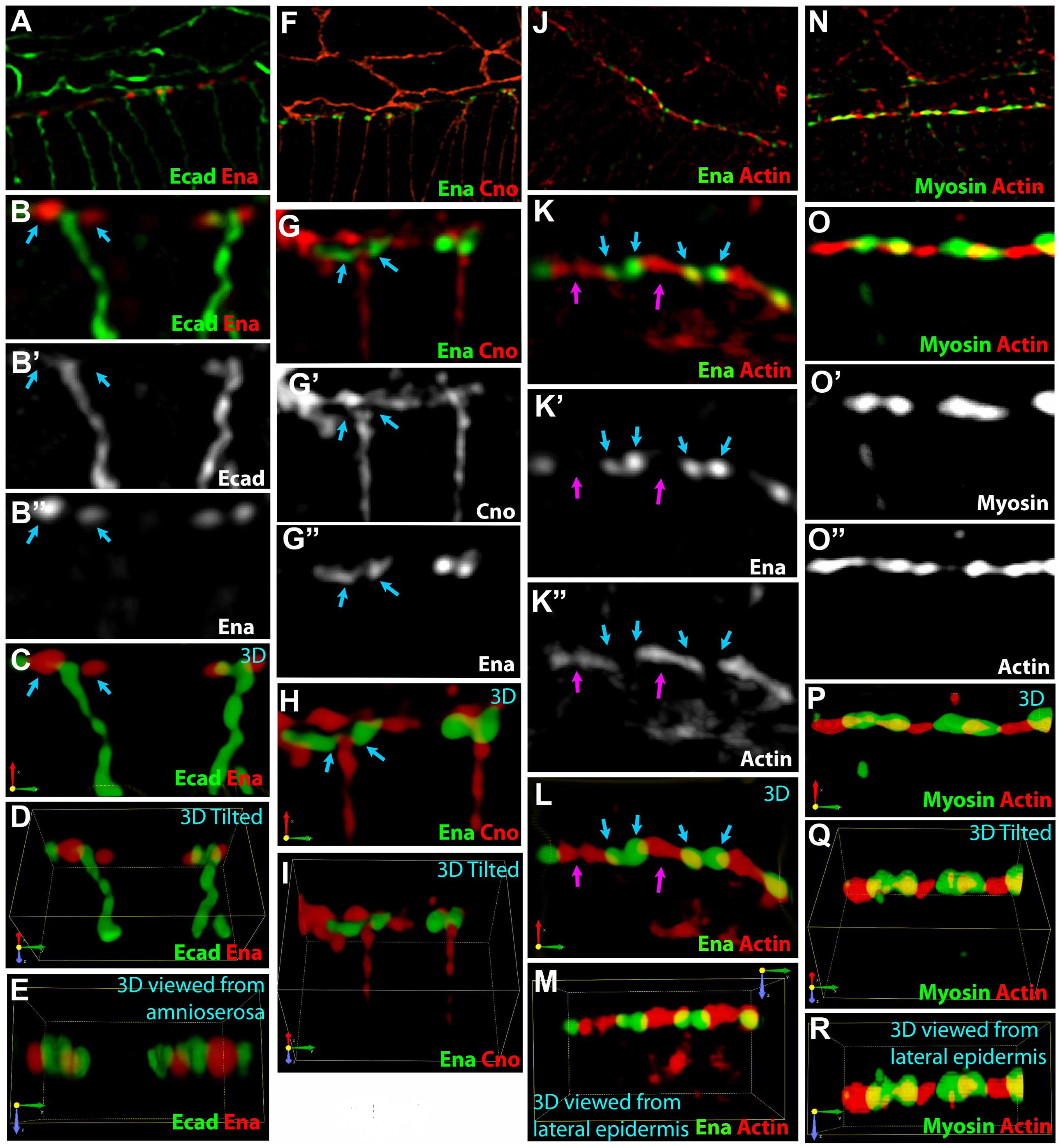
SIM super-resolution microscopy of the LE actin cable and tricellular junctions. SIM images of wild type embryos at mid-stage 13 (mid-dorsal closure), antigens indicated. Directional arrows indicate X (red), Y (green), and Z (blue) axes. A-E. Ena and Ecad. A. Ena puncta (e.g. arrows) flank Ecad at LE tricellular junctions. B-B”. Two tri-cell junctions at the leading edge (LE). SIM resolves two separate Ena puncta (e.g. arrows) that are juxtaposed and surround Ecad at tricellular junctions. Maximum intensity projection (M.I.P.) of z-stacks. C. 3D reconstruction of M.I.P from panel B, using alpha blending. This allows visualization of the leading edge three-dimensional space. D. Tilted view of 3D reconstruction. E. 3D reconstruction from the vantage point of the amnioserosa (rotated 90° from panel C; apical up). Ena and Ecad are in the same plane in apical-basal axis. F-I. Cno and Ena. F. Cno’s relationship to Ena parallels that of Ena and Ecad. G-G” Ena is juxtaposed to Cno at tricellular junctions. H. 3D reconstruction I. Tilted view of 3D. J-M. Ena and actin (visualized with phalloidin). J. Lower magnification view of the LE. K-K”. SIM resolves Ena localization to two separate puncta (blue arrows) located at the ends of each cell’s actin cable (magenta arrows). L. 3D reconstruction. M. 3D reconstruction showing view from the lateral epidermis (rotated 90° from panel L; apical up). Ena and Actin are parallel along the apical-basal axis. N-R. Zipper (Myosin II heavy chain) and Actin. N. Actin and Myosin are arranged in an alternating pattern across the LE of the lateral epidermis. O-O”. Two LE cells. Myosin is enriched in the central portion of each cell’s actin cable. P. 3D view. Q. Tilted view of 3D reconstruction. R. 3D reconstruction side view from the lateral epidermis (rotated 90° from panel P; apical up). Myosin and Actin are parallel along the apical-basal axis. Scale bars in A, F, J & N=20 µm. All others=2 µm.

These data informed our analysis of the cell biological consequences of strong Cno reduction on cell shape, the actomyosin cable and its attachment to the LE cell junctions. Assembling the LE cable puts the LE under tension due to cable contractility (e.g., Kiehart *et al*., 2000; Jacinto *et al*., 2002), with cells exerting force on their neighbors along the cable. In wildtype, this tension is balanced, as assessed by the relatively uniform cell width of different cells along the LE (Fig. 2I, K, red arrows). There is some variability, but it is largely confined to cells near the former segmental grooves (Fig. 2I, blue arrows). In contrast, in most *cnoM-RNAi* embryos, LE cell width is <u>much</u> more variable, with the LE of some cells hyperconstricted (Fig. 2J,L red arrows) while in other cells the LE is hyper-elongated (Fig. 2J,L green arrows). The degree of cell elongation along the dorsal-ventral axis was also much more variable, and some cells failed to elongate (Fig. 2J,L yellow arrows). This data and previous studies of Afadin knockdown in MDCK cells under tension (Choi *et al*., 2016) suggest that Cno/Afadin may help ensure balanced contractility on different cell borders.

We next examined whether cells retained the ability to assemble a planar-polarized actin cable after reduction of Cno. Strikingly, even the most severely affected *cnoM-RNAi* embryos retained at least a partial LE actin cable (Fig. 3H, arrows). In less affected embryos (presumably those inheriting a single copy of the shRNA) the actin cable appeared relatively normal. In embryos with more dramatically altered LE cell shapes, the cable was discontinuous (Fig. 3I, red arrows), though surprisingly even some cells in which the LE was very splayed open retained an actin cable (Fig. 3I-I’, blue arrows). When the LE cells separated from the amnioserosa the cable could remain intact (Fig. 3H, red arrow), but more often the LE cell itself seemed to be ripped apart (Fig. 3J, green and yellow arrows). Loss of AJs could provide a possible mechanism behind cable discontinuity; however, after *cnoM-RNAi* Ecad levels were not substantially lower than wildtype (Fig. 2I vs. J, K vs. L; embryos stained and imaged on same slide, with wildtype marked with histone-GFP), and Ecad continued to be enriched in many LE tricellular junctions (Fig. 3G). However, where contact with the amnioserosa was disrupted Ecad localization was also disrupted (Fig. 2L, blue arrow, 3J). *cnoM-RNAi* embryos also retained junctional localization of the AJ protein Pyd (Fig. 2M vs N). Thus, Cno is not essential for actin cable assembly or contractility, but it is important for maintaining balanced cell contractility and thus actin cable continuity.

Finally, we examined localization of Ena. Ena is also important for maintaining balanced contractility along the LE and for timely and accurate dorsal closure (Gates *et al*., 2007)—the effects of Ena loss on LE cell shapes are quite similar to those seen after *cnoM-RNAi*. Our previous analysis of *cno* zygotic and *pyd* maternal/zygotic mutants revealed alterations in Ena localization and also revealed genetic interactions between *cno* or *pyd* and *ena* (Choi *et al*., 2011). Our stronger knockdown tools allowed us to extend this. Early in dorsal closure *cnoM-RNAi* embryos retained Ena enrichment at lateral epidermal tricellular junctions (e.g., Fig. 4E-E’, yellow arrow) and in segmental groove cells (Fig. 4E’, red arrow) but LE enrichment was reduced (Fig. 4E’, blue arrows). As closure proceeded Ena enrichment became less focused at LE tricellular junctions than in wildtype (Fig. 4F-F”, arrows). In the most severely affected embryos Ena localization became extremely irregular. While occasional tricellular junctions had focused Ena localization (Fig. 4G,H yellow arrows), Ena was not focused at most tricellular junctions (Fig. 4G,H red arrows) and was strongly reduced at others (Fig. 4G, blue arrows). Taken together, our data are consistent with the idea that Cno is important for allowing cell junctions along the LE to resist the contractile force of the actomyosin cable and maintain relatively uniform cell shapes, a role Afadin also plays in MDCK cells (Choi *et al*., 2016). Our data are consistent with the idea that Ena may also play a role in this, via its position at the junction of the cadherin-catenin complex and the actin cable.

### Canoe is critical for cells to retain columnar architecture when challenged by cell division and neuroblast invagination

When we first initiated analysis of Cno a decade ago, we expected it would play an essential role in cell adhesion, with loss leading to complete disruption of epithelial integrity at gastrulation onset, as is seen in mutants lacking Ecad, β- or α-catenin (Cox *et al*., 1996; Tepass *et al*., 1996). However, while many morphogenetic movements of gastrulation are disrupted by Cno loss, the ectoderm initially remains intact (Sawyer *et al*., 2009; Sawyer *et al*., 2011). Cuticle analysis (Fig. 1) suggests that ultimately epidermal integrity is reduced but not eliminated, though the underlying mechanisms remained unexamined. We thus set out to determine what role Cno plays in maintaining epithelial integrity and what mechanisms act in parallel when Cno is absent. To do so, we used our strongest *cno* RNAi condition, *cnoS-RNAi* (the crosses and progeny are diagrammed in Suppl. Fig. 3), which phenocopies complete loss of Cno and leads to very strong reduction in levels of Cno protein (Suppl. Fig. 1A-C). Cuticle analysis revealed results similar to those seen in embryos maternally and zygotically null for Cno (Fig. 1K; Table 1; (Sawyer *et al*., 2009). Head involution and dorsal closure failed, and the thoracic and abdominal epidermis had frequent ventral holes.

We thus examined the progression in epithelial integrity from gastrulation onset to the end of dorsal closure, with a focus on the thorax and abdomen. *cnoS-RNAi* embryos phenocopied *cnoMZ* mutants (Sawyer *et al*., 2009; Choi *et al*., 2013), cellularizing despite disruption in initial establishment of apical AJs and beginning gastrulation with an intact ectoderm (Suppl. Fig. 4A vs B). However, mesoderm invagination was partially to completely disrupted and extension of the germband was slowed or terminated prematurely (Fig. 6F; data not shown). In a subset of embryos (18%; 8/45), the continued intercalation of ectodermal cells combined with the failure to fully extend the germband led to a twisted gastrulation phenotype. In wildtype embryos, germband extension is driven in part by the planar polarized distribution of junctional and cytoskeletal proteins (reviewed in Vichas and Zallen, 2011; Harris, 2018). Actin and myosin are enriched on anterior-posterior (AP) borders, while the cadherin-catenin complex and especially Bazooka (Baz; the fly Par3) are enriched in dorsal-ventral (DV) borders (Suppl. Fig. 4G, yellow versus blue arrows). In *cnoS-RNAi* embryos, this planar polarity was accentuated, with most of the ectoderm hyper-planar polarized and cells arrayed in rows along the DV axis (Suppl. Fig. 4B,H). Arm and Baz planar polarity were substantially enhanced (Suppl. Fig. 4H, blue vs yellow arrows), with reduction in their levels and apical cell separation along AP borders (Suppl. Fig. 4H, yellow arrows), as we observed in *cnoMZ* mutants (Sawyer *et al*., 2011). Baz localization along DV borders also became more fragmentary rather than continuous. Cortical myosin, which is normally planar-polarized to AP boundaries (Suppl. Fig. 4C, I inset), pulled away from the AP cell cortex and cortical levels appeared enhanced (Suppl. Fig. 4D,I, blue arrows vs. I inset). Separation of myosin from the cortex also occurred at multicellular junctions at the center of rosettes (Suppl. Fig. 4C, I, yellow arrows)—intriguingly, tricellular and multicellular junctions were also the points most sensitive to Afadin knockdown in ZO knockdown MDCK cells (Choi *et al*., 2016).

**Figure 6.**
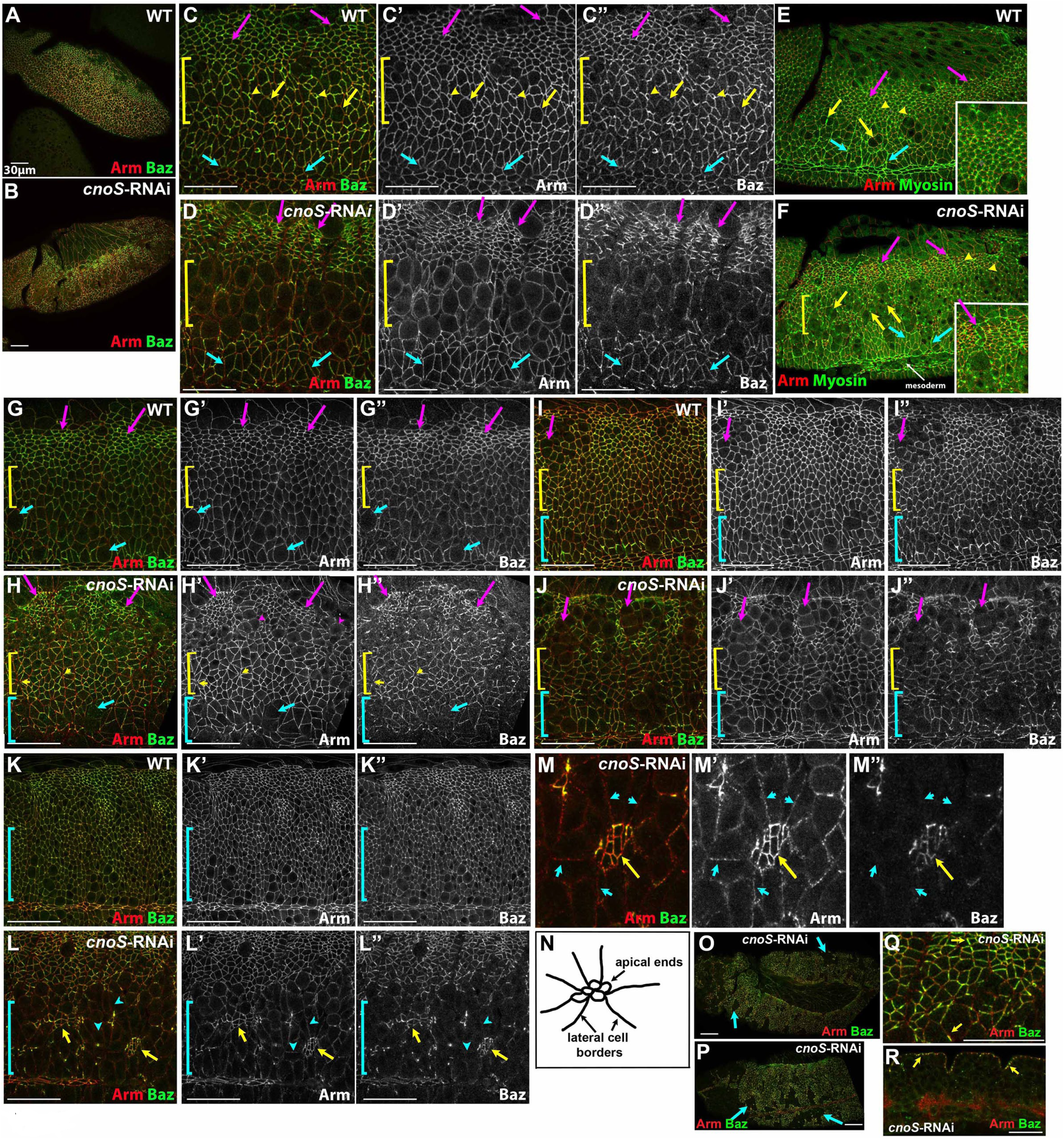
Cno is critical for cells to retain columnar architecture when challenged by cell division and neuroblast invagination. Wildtype and *cnoS-RNAi* embryos, genotypes and antigens indicated. A-F. Stage 9 embryos. A,C,E. Wildtype. Dorsal ectodermal cells (magenta arrows) have competed their first mitotic divisions, resumed columnar architecture and are apically constricted. The bright myosin dots in these cells (E) are midbody remnants of their first mitotic division. Dorsal neurectodermal cells (mitotic domain N) round up individually for mitosis (C, yellow arrows), with reduced cortical Arm and especially Baz, and then resume columnar architecture rapidly (C, yellow arrowheads). Myosin marks contractile rings of dividing cells (E, yellow arrows). Ventral neurectodermal cells (blue arrows), which have not yet divided, have intermediate levels of Baz relative to their more dorsal neighbors. B,D,F. *cnoS-RNAi.* Dorsal ectodermal cells (magenta arrows) are apically constricted but retain strongly enhanced planar polarity of Baz. All dorsal neurectodermal cells remain rounded up (D, F yellow brackets), even though only a subset are still in mitosis, as indicated by myosin marking contractile rings (F,yellow arrow). Ventral neurectoderm cells (blue arrows) resemble wildtype. G,H. Early stage 10. G. In wildtype dorsal ectodermal cells (magenta arrows) remain apically constricted relative to their neighbors. Dorsal neurectodermal cells (yellow bracket) have resumed columnar architecture, while ventral neurectodermal cells (blue arrows; mitotic domain M) have begun to enter mitosis individually. H. In *cnoS-RNAi* embryos Baz localization is strongly reduced in both dorsal ectodermal cells, especially those that remain rounded up (magenta arrows and arrowheads), as well as in the ventral neurectoderm (blue bracket, arrows). While the dorsal neurectoderm is less affected (yellow bracket), Baz localization is less continuous and some cells have formed rosettes (yellow arrowheads). I,J. Late stage 10. I. In wildtype a subset of dorsal ectodermal cells have entered mitosis 15 (e.g., magenta arrows), and ventral neurectoderm cells (blue arrows; mitotic domain M) continue to divide. J. In *cnoS-RNAi* embryos many more dorsal ectodermal (magenta arrows) and ventral neurectoderm cells (blue bracket) remain rounded up with reduced cortical Baz. K,L. Early stage 11. K. In wildtype occasional cells in the neurectoderm (blue bracket) continue to divide individually. L. In more severely affected *cnoS-RNAi* embryos, most cells in the neuroectoderm have lost columnar shape (blue bracket). Islands of cells remain columnar (yellow arrows). These cells are apically constricted and retain elevated cortical Arm and Baz. They appear to form rosettes with lateral cell borders (blue arrowheads) pointing toward the constricted apices. M. Closeup of an “epithelial island” in L. N. Diagram illustrating our interpretation of the “epithelial islands”. O-R. Stage 11 (O,Q) or stage 13 (P) *cnoS-RNAi* embryos highlighting epithelial holes (blue arrows) and unevenness of Baz accumulation, even in regions of where the epidermis remains columnar (Q, yellow arrows). R. Cross section through the epidermis of a stage 13 embryo. Baz and Arm remain apically polarized (arrows). Scale bars=30µm.

Maintaining integrity of the ectoderm and epidermis requires the cells to maintain epithelial architecture while managing the challenge of the substantial AJ remodeling involved in three processes: cell intercalation during germband elongation, cell rounding during mitosis and the subsequent return to a columnar architecture, and, in the neurectoderm, invagination of ~30% of the ectodermal cells as neural stem cells (neuroblasts; NBs). Beginning at the end of stage 7 of embryogenesis, cells across the ectoderm undergo mitosis in programmed groups called mitotic domains (Foe, 1989). As cells enter mitosis, they round up and cortical AJ protein accumulation per unit membrane is reduced (e.g., Suppl. Fig. 4E, arrows), as was previously observed in the pupal notum (Pinheiro *et al*., 2017). NB invagination and the cell rearrangements that drive germband extension overlap with the onset of mitosis of some of the later mitotic domains. This is known to make the ventral epidermis more sensitive to reductions in function in either junctional or apical polarity proteins (Tepass *et al*., 1996; Harris and Tepass, 2008). The trunk ectoderm during stage 9 (Fig. 6A, C) can be roughly divided into three regions: 1) the dorsal ectoderm (Fig. 6C, magenta arrows), in which the cycle 14 mitoses happen relatively early and from which no NBs delaminate, 2) the more dorsal neurectoderm (Fig. 6C, yellow bracket) —mitotic domain N—in which individual cells divide (Fig. 6C, yellow arrows) during embryonic stage 9, and 3) the more ventral neurectoderm (Fig. 6C, blue arrows)-mitotic domain M— in which cells divide during embryonic stage 10. In mitotic domains N and M cells round up and undergo mitosis individually rather than collectively Fig. 6C, yellow arrows), and this occurs while a subset of the cells invaginate to become neuroblasts (Fig. 6C, arrowheads). Mitotic cells then rapidly resume a columnar shape, and thus only a subset of cells are rounded up at any given time. During this process, the apical ends of adjacent dorsal epidermal cells become smaller (Fig. 6C, magenta arrows), potentially because their apical contractility is less restrained by their ventral neighbors.

In *cnoS-RNAi* embryos early mitotic domains appeared to fire roughly on schedule at stages 7 and 8 (Suppl. Fig. 4F vs E., data not shown). However, as germband extension continued and cells in the neuroectoderm initiated mitosis, *cnoS-RNAi* embryos deviated dramatically from wildtype (Fig. 6B). Virtually all cells in the dorsal neurectoderm (mitotic domain N) were simultaneously rounded up, with reduced cortical cadherin (Fig. 6D, bracket), suggesting that they are slower to resume columnar cell shape. Dorsal ectodermal cells became even more highly apically constricted (Fig. 6D, magenta arrows) than their wildtype counterparts (e.g. Fig. 6C), potentially reflecting reduced contractility in their rounded up ventral neighbors. Dorsal ectodermal cells also formed clear rows along the AP axis. At stage 9, cells of the ventral neurectoderm, which have not yet undergone mitosis, were more similar to their wildtype counterparts (Fig. 6D vs. C, blue arrows). Myosin staining highlighted the same differences. In wildtype, cells in the dorsal ectoderm (Fig 6E, magenta arrows) had completed division, with myosin dots marking the remnant midbody (Fig. 6E, arrowheads). Mitotic cells in the dorsal neurectoderm could be identified by their cleavage furrows (Fig. 6E, yellow arrows, while ventral neurectoderm cells retain planar polarized cortical myosin (Fig. 6E, blue arrows). In *cnoS-RNAi* embryos, while dorsal ectodermal cells had completed mitosis, as indicated by remnant midbodies (Fig. 6F, arrowheads), and rows of hyper-planar polarized cells were separated and had elevated cortical myosin (Fig. 6F, magenta arrows). While some rounded up cells in the dorsal neurectoderm were actively dividing, as indicated by their cleavage furrows, others had completed division, as indicated by the midbody staining, but not resumed columnar architecture (Fig. 6F, yellow arrows). In contrast, the ventral neurectoderm (Fig. 6F, blue arrows) remained relatively normal.

These differences continued and became more accentuated during stage 10. In wildtype, cells in the dorsal neurectoderm resumed a columnar architecture (Fig. 6G, bracket), while cells in the ventral neurectoderm (mitotic domain M) entered mitosis individually (Fig. 6G, blue arrows). Slightly later, some cells began their 15^th^ round of mitosis (e.g. Fig. 6I, magenta arrow). In contrast, in *cnoS-RNAi* embryos most cells in the ventral neurectoderm and some in the dorsal neurectoderm remain rounded up with reduced cortical Armadillo (Arm; *Drosophila* β-catenin; Fig. 6H, J, blue bracket and arrows). Cells entering mitosis 15 seemed to also be delayed in resuming columnar shape (Fig. 6J, magenta arrows). Cell separation was observed at multicellular junctions, suggesting they are weak points in a relatively intact epithelium (Fig. 6I, yellow arrows). As *cnoS-RNAi* embryos entered stage 11, the delay in the resumption of columnar shape in the neurectoderm was followed by a loss of epithelial integrity, particularly along the ventral midline (Fig. 6L vs. K, brackets). These epithelial disruptions remained through the end of germband retraction and dorsal closure (Fig. 6O,P arrows) and likely lead to the holes in the ventral cuticle we observed. This data suggests a likely role for Cno in re-establishment of columnar epithelial architecture after the challenges posed by mitosis, NB invagination and cell intercalation.

During establishment of apical-basal polarity, Cno regulates apical positioning of both AJs and the apical polarity determinant Baz during cellularization (Choi *et al*., 2013), and *cno* mutants have accentuated planar polarity of Ecad and especially Baz during germband extension (Sawyer *et al*., 2011). Baz plays a key role in maintaining epithelial integrity (Müller and Wieschaus, 1996). We thus examined the effects of Cno knockdown on Baz localization in the ectoderm when cells were challenged by NB invagination and mitotic divisions. In wildtype Baz localization largely parallels that of Arm. Both Arm and Baz are reduced in cortical intensity in cells rounded up for division, though the reduction is somewhat more pronounced for Baz (Fig. 6C”, yellow arrows). In contrast, by stage 9 *cnoS-RNAi* has much more severe consequences for Baz localization. In the apically constricted cells of the dorsal ectoderm, Baz remains exceptionally planar polarized, being essentially restricted to DV cell borders (Fig. 6D”, magenta arrows). In the rounded-up cells of the dorsal neurectoderm, Baz is almost completely lost from the cell cortex (Fig. 6D”, yellow bracket). These differences from wildtype become even more accentuated in stage 10. In rounded up cells of the neurectoderm (Fig. 6H, blue bracket) and dorsal ectoderm (Fig. 6H, magenta arrows), Baz levels are reduced even more than those of Arm, while Baz levels remain high in the hypercontracted dorsal ectodermal cells (Fig. 6H”, magenta arrows). Even in less disrupted regions of the epidermis Baz localization became more uneven, with some cell borders having much higher levels than others (Fig. 6Q, arrows), although the overall apical basal polarization of Arm and Baz remains relatively normal (Fig. 6R).

By stage 11, epithelial architecture is lost in the ventral neurectoderm, with small “islands” of apically constricted cells seemingly separated from one another (Fig. 6L,M yellow arrows). We suspect this reflects a situation where a subset of cell junctions fail, leading to unbalanced apical contractility and allowing groups of cells to apically constrict when they were released from the contractile restraint of their neighbors. The bright islands of Arm and Baz staining would thus be the constricted apical ends of these cell groups, while the adjacent regions that stain for Arm and not Baz represent the lateral borders of the apically constructed cells in the island (Fig. 6L,M blue arrows; diagrammed in 6N). This would parallel what we observed in Afadin ZO knockdown MDCK cells, with some cell junctions failing and a subset of cell borders hyper-constricted while others were hyper-extended (Choi *et al*., 2016). Together, these data suggest that Cno plays an important role in allowing cells to maintain epithelial integrity when challenged by junctional remodeling during morphogenetic movements, and also suggest that the polarity protein Baz is most sensitive to Cno loss. However, they also reveal that many cells can maintain columnar epithelial architecture even when Cno is lost, in contrast to cells lacking Baz or core AJ proteins.

### Pyd/ZO-1 works in parallel with Canoe to maintain epithelial integrity

Afadin, Cno’s mammalian homolog, acts in parallel with ZO-1 family proteins to maintain epithelial integrity in cultured mammalian cells (Choi *et al*., 2016). In *Drosophila*, unlike in mammals, both Cno and Pyd localize to AJs (Takahashi *et al*., 1998; Wei and Ellis, 2001; Jung *et al*., 2006; Seppa *et al*., 2008; Sawyer *et al*., 2009; Choi *et al*., 2011). Strikingly, Pyd localization mirrors that of the AJ proteins even in the one place where Cno and AJ proteins differ subtly. During germband extension, Baz and AJ proteins are subtly enriched on DV cell borders (Zallen and Wieschaus, 2004; Blankenship *et al*., 2006; Harris and Peifer, 2007)--this is more apparent for Baz, whose planar polarization is more pronounced (Suppl. Fig 5A”, yellow versus blue arrows.) At this stage, we found Pyd localizes like the other AJ proteins (Suppl. Fig 5A’” vs A’). In contrast, Cno is enriched on AP cell borders (Sawyer *et al*., 2011), like myosin and actin (Bertet *et al*., 2004; Zallen and Wieschaus, 2004). Loss of Cno enhances the planar polarity of Arm and Baz, by reducing their levels on AP cell borders (Sawyer *et al*., 2011); Suppl. Fig 5B’=B”, yellow vs. blue arrows), and in parallel enhances the planar polarity of Pyd (Suppl. Fig 5B”’, yellow vs blue arrows. However, Cno loss does not dramatically alter levels of cortical Pyd (Sawyer *et al*., 2009; Suppl. Fig. 5B vs. A, D vs C).

Cno and the ZO-1 homolog Pyd interact both physically and genetically (Takahashi *et al*., 1998), supporting the idea that they might act together or in parallel. However, previous studies of genetic interactions were confined to partially functional alleles. To determine whether and how Cno and Pyd acted in parallel we sought to combine strong reduction of both. The close proximity of the two genes in the genome precluded combining null alleles of both on the same chromosome by recombination. We chose a different approach, combining complete maternal and zygotic loss of Pyd (Choi *et al*., 2011) with strong (though not complete) maternal and zygotic reduction of Cno via RNAi (*matGAL4/+; cnov20shRNA pyd/pyd* females × *cnov20shRNA pyd/+* males; heterozygous males were chosen due to the semi-sterility of *pyd* homozygous males; crosses and resulting progeny are diagrammed in Suppl. Fig. 3). We can distinguish by immunofluorescence analysis the 50% that are maternally <u>and</u> zygotically *pyd* mutant by their lack of Pyd staining, and the strongest 25% of the embryos will also have two zygotic copies of the *cnov20shRNA* (*cnov20shRNA pyd/ cnov20shRNA pyd*— referred to as *pyd cno* double mutants below). As a control for Cno reduction alone, we compared our double mutants to embryos with the same degree of Cno reduction but retaining wildtype levels of Pyd (*matGAL4/+; cnov20shRNA /+* females x *cnov20shRNA /+* males). Cell biological analysis revealed that the more severe embryos from this control cross (likely those receiving two copies of the *cnov20shRNA* zygotically) were similar in phenotype to those of *cnoS-RNAi* embryos (Suppl. Fig. 6). We also compared the phenotype of double mutants to our strongest *cno* RNAi (*cnoS-RNAi*, the genotype analyzed above), which immunoblotting (Suppl. Fig. 1) and cuticle analysis (Fig. 1) suggest phenocopy complete loss of Cno.

To confirm the genotypes and assess the degree of Cno knockdown, we performed immunoblotting. The combination of GAL4 driver and *cnov20shRNA* used in our control cross produced nearly complete knockdown of maternal Cno, as assessed in 1-4 hr old embryos (3.6% of wildtype; Suppl. Fig. 1D-E). Knockdown remain strong but not complete at 12-15 hours (8.1% of wildtype; Suppl. Fig 1D,F)—due to variability in the number of *cnov20shRNA* copies they receive, late Cno knockdown likely varies among embryos. Embryos from the *pyd cno* cross had similar levels of Cno knockdown (2.7% of wildtype at 1-4 hrs and 6.5% of wildtype at 12-15 hrs; Suppl. Fig. 1D-F; we also verified knockdown by immunofluorescence; Suppl. Fig. 1J vs I). Our analysis also verified the strong reduction of Pyd in this population (Suppl. Fig. 1D-F)—the remaining Pyd seen in 1-4 hr embryos and the partial return of Pyd at 12-15 hrs reflect the 50% of embryos that are zygotically rescued. We also examined effects on accumulation of the AJ protein Arm. Arm levels were essentially unchanged at 1-4 hrs and somewhat reduced at 12-15 hrs (Suppl. Fig. 1), consistent with our earlier analysis of *cnoMZ* mutants (Sawyer *et al*., 2009).

As an initial analysis of the effects of reducing both Pyd and Cno on morphogenesis, we examined effects on morphogenesis and epithelial integrity by analyzing embryonic lethality and cuticle patterning. The Cno knockdown cross that serves as the control for our double mutant (cross and progeny diagrammed in Suppl. Fig. 3) led to partially penetrant embryonic lethality (84% lethality, n=696; Fig. 7J). Among the lethal embryos, many had a wildtype or only mildly defective cuticle (21%; Fig. 7A-I,K, Table 1; these embryos and those that are embryonically viable are likely the 25% carrying no zygotic copy of the *cno20shRNA* construct). Most of the rest had significant defects in head involution and dorsal closure, with the most severe also having holes in the ventral epidermis (roughly 10%; Fig. 7K, Table 1). Pyd loss alone has milder effects, even in maternal zygotic null mutants, with 60% embryonic lethality and only mild defects in head involution and dorsal closure in the non-viable embryos (Choi *et al*., 2011). In contrast, embryos from the *pyd cno* double mutant cross were 100% lethal (n=374; Fig. 7J) and exhibited much more severe defects in morphogenesis and epithelial integrity (Fig. 7K,L; Table 1). In addition to complete failure of head involution and dorsal closure, 69% had defects in epidermal integrity. Most striking, 24% had only pieces of intact cuticle remaining, a fraction equivalent to the quarter of the progeny that were both *pyd* maternal and zygotic mutant and had the strongest *cno* knockdown. Importantly, the defects of those embryos were significantly more severe than those of *cnoS-RNAi* embryos (Fig. 7K,L; Table 1), which have essentially complete *cno* maternal/zygotic knockdown (Suppl. Fig. 1). These data suggest that Cno and Pyd act in parallel to ensure epithelial integrity.

**Figure 7.**
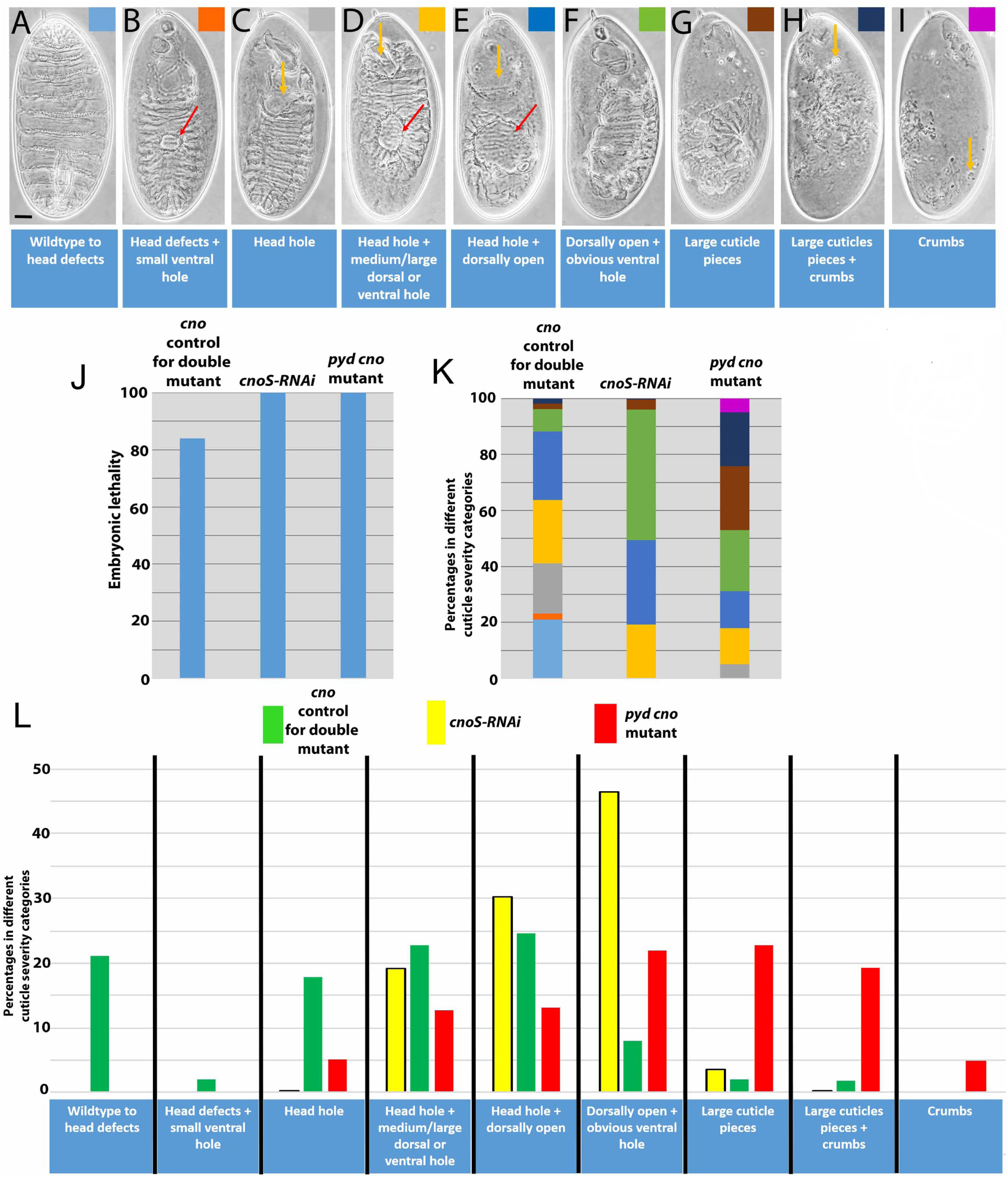
Loss of Pyd substantially enhances the effects of Cno loss on morphogenesis and epithelial integrity. A-I. Cuticle preparations revealing the spectrum of defects in morphogenesis and epithelial integrity seen in *cnoS-RNAi, pyd cno*, and *cno control* knockdown embryos, as in Figure 1. Scale bar= 30µm. J. Embryonic lethality. K,L. Two illustrations of the enhancement of the severity of the Cno knockdown cuticle phenotype by loss of Pyd. *pyd cno* mutants have defects that are much more severe than the corresponding *cno RNAi control* and that are even more severe than those of *cnoS-RNAi*.

### *pyd cno* mutants exhibit defects in epithelial integrity from the onset of gastrulation, and tricellular and multicellular junctions are the weakest points

We thus explored the cell biological basis of these strong defects in epithelial integrity. While the *pyd cno* cross produces embryos of four genotypes (Suppl. Fig. 3), by staining embryos with Pyd antibody we could focus on the half that were *pyd* maternal/zygotic mutant, thus lacking Pyd staining. All had equivalent maternal Cno knockdown, and they varied in the amount of zygotic Cno knockdown. Below we refer to these as *pyd cno* mutants. During gastrulation onset, they shared with *cnoMZ* and *cnoS-RNAi* mutants defects in apical positioning of AJs and Baz during cellularization (Suppl. Fig. 7A; Choi *et al*., 2013), defects in mesoderm invagination (Suppl. Fig. 7B,C), strongly accentuated planar polarity of AJs and Baz (Suppl. Fig. 7C, C inset, magenta arrows), and enhancement of cortical myosin along with myosin disconnection from cell junctions at AP cell borders as germband extension initiated (Suppl. Fig. 7D,E). The hyper-planar polarization of Baz and Arm at DV borders and myosin at AP borders, as well as cell alignment into rows (Suppl. Fig. 7C inset, yellow arrows) were further accentuated relative to *cnoS-RNAi* embryos, with grooves appearing along the AP axis between adjacent rows of cells (Suppl. Fig. 7C inset, blue arrows). Two particularly deep grooves appeared to coincide with the positions of the normally transient dorsal folds (Suppl. Fig. 7C, yellow arrows). The first mitotic domains, including mitotic domain 11 in the trunk, fired in a timely manner (Fig. 8A,C vs B,D, white arrows), but cortical Arm in dividing cells was less continuous and large gaps appeared at multicellular junctions between some rounded up cells (Fig. 8C vs D, blue arrows). In *pyd cno* mutants adjacent non-dividing cells in the neurectoderm formed rosettes with gaps in the center at multicellular junctions (Fig. 8E,F, magenta arrows). F-actin staining also highlighted the gaps at tricellular/multicellular junctions between dividing cells (Fig. 8G, H, blue arrows) and at the center of rosettes of non-dividing cells at multicellular junctions (Fig. 8H, magenta arrows). In *pyd cno* mutants cortical myosin levels were highly elevated at AP cell borders (Fig. 8I,J) and myosin separated from the cortex at many vertices (Fig. 8J, arrows). These data suggest *pyd cno* mutants are even more sensitive to the stresses of rounding up for division and of cell rearrangements during germband extension, that tricellular and multicellular junctions are the weakest points, and that loss of both Pyd and Cno strongly accentuates the reciprocal planar polarity of Baz/Arm versus myosin.

**Figure 8.**
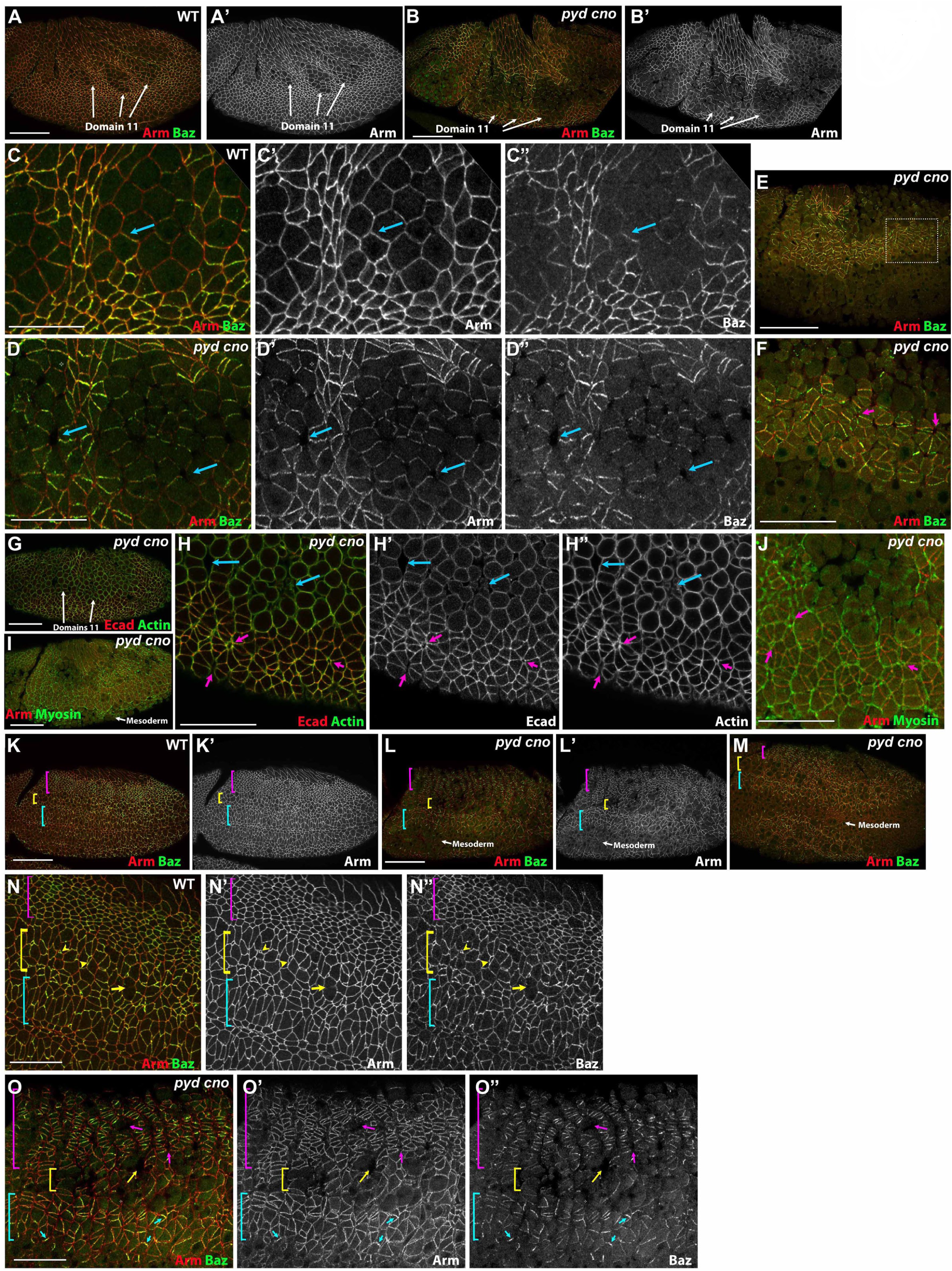
Loss of epithelial integrity begins earlier in *pyd cno* double mutants and Baz localization is affected earlier than that of AJs. A-J. Stage 8 embryos focused on early mitotic domain 11. A,B. This mitotic domain fires on time in *pyd cno* mutants. A-D. Cortical Arm and Baz accumulation are reduced in dividing cells in both wildtype (A,C, arrows) and *pyd cno* (B,D, arrows). However, in *pyd cno* mutants large gaps appear between dividing cells (D, arrows) that are not seen in wildtype. E,F. In *pyd cno* mutants, in more ventral cells that have yet to divide both Arm and Baz become hyper-planar polarized and gaps appear at multicellular junctions where cells are forming rosettes (F, arrows). Dotted box in E indicates area enlarged in panel F. G,H. In *pyd cno* mutants staining with Ecad and actin also highlights large gaps between dividing cells in mitotic domain 11 (H, blue arrows) and gaps at the center of rosettes more ventrally (H, magenta arrows). I,J. In *pyd cno* mutants cortical myosin is elevated at anterior-posterior boundaries and often separates from the cortex, particularly at tricellular and multicellular junctions (J, magenta arrows). K-O. Stage 9 embryos. K, N. In wildtype dorsal ectodermal cells (magenta brackets) have completed their first division, apically constricted and resumed columnar shape. Some cells of the dorsal neurectoderm (yellow brackets) are individually dividing (N, yellow arrow) while others invaginate as neuroblasts (N, yellow arrowheads). Cells of the ventral neurectoderm (blue brackets) have yet to divide. L,M,O. *pyd cno* mutants. Dorsal ectoderm cells (magenta brackets) form prominent rows along the dorsal ventral axis, with hyper-planar polarized Arm and Baz. Gaps appear between many of the rows (O, magenta arrows). Dividing cells of the dorsal neurectoderm (yellow brackets) have reduced Arm and Baz staining and gaps appear between cells (O, yellow arrow). In the ventral neurectoderm (blue brackets), while Arm cortical accumulation remains largely intact, Baz cortical localization has become fragmented, with only some cell borders retaining Baz (O,blue arrows). Scale bars in A,B,E,G,I E,K & L =20 µm. All other scale bars=10µm

### *pyd cno* mutants exhibit loss of cortical Bazooka localization preceding widespread loss of epithelial integrity

*pyd cno* mutants roughly resembled *cnoS-RNAi* embryos as cells in the dorsal neuroectoderm (mitotic domain N) began to round up for division (Fig. 8K vs L,M vs Fig. 6D), possessing hyperconstricted dorsal ectoderm cells arrayed in rows (Fig. 8N vs O, magenta brackets) and virtually all cells in the dorsal neurectoderm rounded up (Fig. 8N vs O, yellow brackets). However, the ectoderm was more significantly disrupted than in *cnoS-RNAi*. As cells separated apically, grooves along the AP axis between rows of dorsal ectodermal cells were further attenuated and holes appeared (Fig. 8O, magenta arrows). Large gaps appeared between dividing cells in the dorsal neurectoderm (Fig. 8O, yellow arrow). In the ventral neurectoderm (Fig. 8N vs O, blue brackets), junctional accumulation of Baz became widely fragmented in *pyd cno* mutants (Fig. 8O, blue arrows; 38% of *cno pyd* mutants have these stronger phenotypes at stage 9, n=19), in contrast to the more intact Baz staining seen in *cnoS-RNAi* embryos (Fig. 6D). Thus, loss of Pyd continued to strongly enhance the effects on epithelial integrity of Cno knockdown.

As embryos proceeded into stage 10, loss of epithelial integrity *pyd* cno mutants became even more widespread (Fig. 9A vs B,C), ranging from embryos at the most severe end of the phenotypes seen in *cno*S-RNAi embryos (Fig. 9B), to those with global disruption of the epidermis (Fig. 9C). At this stage in wildtype, most cells across the epidermis have resumed a columnar architecture (Fig. 9D), with small groups of cells or individual cells in mitosis 15 (Fig. 9D, arrow). In contrast, in most *pyd cno* mutants (55% of *pyd cno* mutants at stage 10; n=50), cells of the neurectoderm formed into separate “balls” (Fig. 9E, white arrows) segregating from their neighbors, and as a result the surface of the embryo became highly irregular. In these regions, AJ proteins and Baz staining became highly fragmented (Fig. 9E, blue arrows), with only small groups of cells retaining cortical Arm. As we saw at stage 9, junctional accumulation of Baz was much more strongly affected than that of Arm (Fig. 9E, yellow and magenta arrows). Strikingly, the amnioserosa was much less affected (Fig. 9B,C, red arrows).

**Figure 9.**
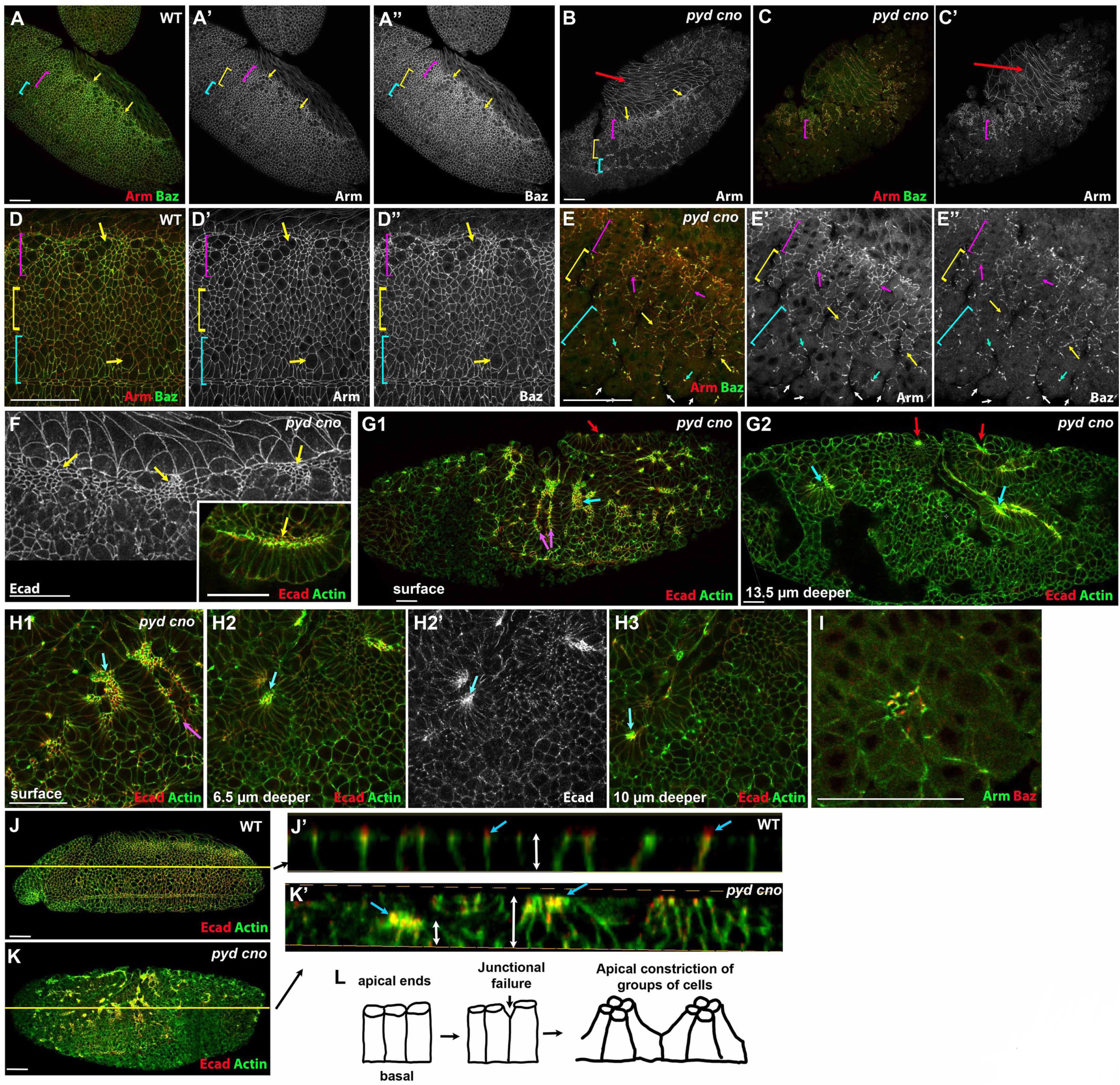
In *pyd cno* mutants epidermal integrity is rapidly lost as cells separate into epithelial islands and balls. A-E. Late stage 9/early stage 10 embryos. A,D. Wildtype. Dorsal ectoderm cells (D, magenta bracket) are generally columnar and apically constricted with a few entering mitosis 15 (D, top arrow). Dorsal neurectoderm cells (yellow bracket) have largely completed division and resumed columnar architecture. Individual cells of the ventral neurectoderm (blue bracket) continue to divide (D, bottom arrow). B,C,E. In *pyd cno* mutants epidermal integrity is rapidly lost. B. Less severe example. Dorsal ectoderm (arrows) and some dorsal neurectodermal cells (magenta bracket) remain epithelial while only small epithelial islands remain in the ventral neurectoderm (blue bracket). C,E. More severely affected embryo. At the surface, only small groups of cells in the dorsal ectoderm (E, magenta arrows) and dorsal neurectodermal cells (E, yellow arrows) remain epithelial, and even in these cells cortical Arm localization is more intact than is Baz. In the ventral neurectoderm only junctional fragments remain (E, blue arrows). F,F inset. In less severely affected *pyd cno* embryos, dorsal ectodermal cells become highly apically constricted (arrows). This often results in groups of ectodermal cells at the border of the amnioserosa folding inward, with their apical ends turned ventrally (inset). G,H. Widespread loss of epithelial integrity in *pyd cno* mutants fragments the embryo into groups of cells that remain associated with one another but which have lost junctional contact with other neighbors. Groups of cells fold inward to form epithelial folds and balls. This is readily seen by taking different sections through an embryo. G1, H1. Surface views. Only small patches of cells retain columnar architecture, and these have apical ends that are severely apically constricted (blue arrows). Other cells can be seen folding inward as sheets (magenta arrows) or rosettes. G2,H2,H3. Deeper sections through the embryo reveal other groups of cells that have been internalized as epithelial balls (G2, red arrows), rosettes or sheets (blue arrows). In these Ecad and actin remain enriched apically. I. In some epithelial balls Baz can also be observed apically retained. J-K. Stage 10 embryos. J,K. Surface views. Lines indicate plane of section shown in closeups. J’,K’. Cross sections generated using Imaris. J’. In wildtype cells are even in height (double-headed arrow), columnar in architecture, and have Ecad enriched at the apical junctions (blue arrows). K’. In *pyd cno* mutants the embryo surface becomes very uneven, with cells varying in height (double-headed arrows). Ecad and actin remain enriched at apical ends of groups of apically constricted cells (blue arrows). L. Diagram illustrating our interpretation of the progression in *pyd cno* mutants. Cells begin columnar (left). A subset of cell junctions fail (middle) and groups of cells then apically constrict (right). Scale bars=20µm in panels A-E, G1-G2. All other scale bars=10µm.

In *pyd cno* mutants, unbalanced contractility was even more pronounced than after *cno*S*-RNAi*, as groups of cells separated from their neighbors. The enhanced apical constriction of dorsal ectodermal cells led to the folding of these cells inward at the junction with the amnioserosa (Fig. 9F, F inset). Meanwhile small islands of epithelial cells in the neuroectoderm formed epithelial balls and sheets. Many of these became internalized into the embryo, as was revealed by imaging successively deeper sections into the embryo (Fig. 9G1 vs G2, H1 vs H2,H3, arrows indicate groups of apically constricted cells). However, in these epithelial cell clusters Ecad, actin (Fig. 9H, arrows), and occasionally Baz (Fig. 9I) remained enriched on their apical ends. The three-dimensional nature of these epithelial disruptions was reinforced when we used Imaris to generate 3D renderings of the surface of wildtype and mutant embryos (Fig 9J,K). Cross-section slices through wildtype embryos revealed cells of uniform heights and columnar architecture, with apical enrichment of Ecad (Fig. 9J’). In contrast, sections through *pyd cno* mutants revealed separate groups of apically constricted cells with variable heights (Fig. 9K’). These data suggest that in *pyd cno* mutants the junctional remodeling accompanying mitosis and neuroblast invagination imposes stress on adhesion leading to separation at weaker cell interfaces (diagrammed in Fig. 9L). Groups of cells that retain cell adhesion then apically constrict and are internalized as large folds or epithelial balls. Together, these data reveal that Pyd and Cno work in parallel to maintain epithelial integrity when challenged by the junctional remodeling inherent in morphogenesis, and further suggest that they may exert their effects by regulating Baz cortical localization during this process.

## Discussion

Linkage of the actomyosin cytoskeleton to cell-cell and cell matrix junctions drives cell shape change during normal embryonic development and adult homeostasis. One key question for the field involves defining how cell-cell junctions are dynamically regulated to allow cells to move within the plane of the epithelium and change shape without disrupting epithelial integrity. The *Drosophila* embryo provides a superb place to test hypotheses, as cells divide, ingress, and undergo the shape changes and cell rearrangements involved in morphogenesis. Here we explored how the junctional proteins Cno/Afadin and Pyd/ZO-1 act together or in parallel to maintain epithelial integrity.

### Cno: more than just a junction:cytoskeletal linker

We initially pursued Cno as a potential core component of the AJ, like Ecad and the catenins. However, loss of Cno does not lead to immediate loss of epithelial architecture; instead, it disrupts a series of cell shape changes requiring linkage of the actomyosin cytoskeleton to AJs (Sawyer *et al*., 2009; Sawyer *et al*., 2011). This led us to hypothesize that Cno acts as one of several proteins that link junctions and the cytoskeleton, and that it reinforces these linkages under mechanical tension. Our work on Afadin in mammalian MDCK cells (Choi *et al*., 2016) suggested a further modification, as loss of Afadin did not uniformly disconnect junctions from the cytoskeleton. Instead, tricellular and multicellular junctions were most sensitive to Afadin loss. Our data suggested this resulted from those junctions being the site of end-on linkage of bicellular actin cables to cadherin-based junctions, and thus the place with the highest “molecular tension” on junctional proteins. Afadin knockdown disrupted junctional-cytoskeletal linkage at tricellular junctions, and the normally tightly bundled junctional actin and myosin spread broadly across the lateral membrane. These local disruptions could then spread to adjacent bicellular junctions. The MDCK cell data also suggested that reducing Afadin function led to loss of contractile homeostasis, with some cell borders becoming hyper-constricted and others hyper-extended. This prompted us to return to the embryo, with these insights in mind.

Our current data support and amplify on this idea. During dorsal closure, the leading edge (LE) provides an interesting model of balanced contractility. The planar-polarized supercellular actomyosin cable along the LE provides part of the force-generating machinery that powers dorsal closure (reviewed in Hayes and Solon, 2017; Kiehart *et al*., 2017; though see Ducuing and Vincent, 2016; Pasakarnis *et al*., 2016). Each border is a separate contractile unit, pulling on its neighbors, but in the wildtype state each cell’s cable is equivalently contractile relative to its neighbors, maintaining relatively uniform LE cell widths across the cable. Our SIM super-resolution imaging builds on earlier work, supporting the idea that the LE actin cable is linked cell-cell at each LE AJ, with Ecad and Cno enriched at LE tricellular junctions, and Ena localization suggesting actin barbed ends are also enriched at those sites. Myosin is enriched at the medial regions of each LE cable. We found that strong reduction of Cno function does not abrogate actin cable assembly at the LE, nor does it block contractility. Instead, LE contractility appears to become unbalanced, with LE cells becoming hyper-constricted or hyper-elongated. This occurs in parallel with disruption of actin architecture at the LE, as reflected by loss of focused Ena localization at LE junctions, which may reflect alterations in the localization of actin barbed ends. We speculate that this reflects failure of end-on actin attachment at a subset of LE tricellular junctions. These cells then cannot counter the contractile force of their neighbors, and splay open, while their neighboring cells, released from the resistance of those neighbors, hyper-constrict. These data are quite consistent with the junctional failure at tricellular and multicellular junctions and the unbalanced contractility we observed in MDCK cells after Afadin ZO knockdown (Choi *et al*., 2016).

The *pyd cno* mutant phenotypes also are interesting when viewed from this perspective. In mutants cells form rows and rosettes from gastrulation onset, with the cell separation and loss of junctional integrity focused at AP cell borders and multicellular junctions. Later on this progresses to more global epithelial disruption. One speculative interpretation of these disruptions is that, as we observed in Afadin ZO knockdown MDCK cells, junctional integrity and junction-cytoskeletal linkage failure first occurs at tricellular junctions, but after the initial disruption spreads more broadly. This leads to unbalanced contractility between different groups of cells. Cells that retain junctional connections apically constrict, as they are no longer effectively resisted by adjacent cells in which junctional connections have been weakened or lost. Strikingly, in *pyd cno* mutants these groups of cells sometimes go on to move inward, forming epithelial rows or balls that go on to secrete the cuticle pieces we observe in the terminal phenotype.

One place where unbalanced contractility is a necessary feature of normal cell rearrangements is during germband extension. Wildtype embryos rely on “regulated unbalanced contractility”, resulting from the planar polarization of junctional and cytoskeletal proteins (reviewed in Kong *et al*., 2017)). Actin and myosin accumulate at elevated levels on anterior-posterior (AP) borders while adherens junction proteins and Baz are elevated at dorsal-ventral (DV) borders. In wildtype these differences are subtle, but they drive constriction of AP borders and thus cell rearrangements. However, adhesion remains robust enough that this does not lead to disruption of cell adhesion and thus cell separation. Intriguingly, Cno is elevated at AP borders along with actin and myosin, rather than at DV borders with other junctional proteins (Sawyer *et al*., 2011). We speculated it may act there to make sure the connection of AJs to the actomyosin cytoskeleton is not disrupted during contractility. In contrast, in *cno* mutants the planar polarity of AJs and Baz is substantially elevated, due to reduced accumulation on AP borders (Sawyer *et al*., 2011). Our data here suggest cortical myosin is elevated on the opposing AP borders. This leads to preferential cell separation at AP borders, where AJ levels are reduced and contractility is elevated. These differences are even more accentuated in *pyd cno* mutations. This fits well into an emerging idea that cell adhesion and cortical actomyosin are inversely correlated, in a balance between adhesion strength and cortical tension (Maitre and Heisenberg, 2011, 2013; Winklbauer, 2015). Thus, one might speculate that the reduction in adhesion at a particular cell border in *cno* or *pyd cno* mutants might lead to elevated actomyosin contractility, which might in turn further reduce adhesion. This kind of run-away feedback process might explain why adhesion fails at a subset of cell borders, leading to cell separation and the formation of cell groups and epithelial balls.

When speculating about mechanism, it is useful to note what does and does not go wrong after Cno loss or in *pyd cno* mutants. Even in severely disrupted regions of the embryo, cells that retain junctional connections also retain epithelial polarity, as reflected by continued apical junctional enrichment of Ecad, Arm, and Baz and the apical enrichment of actin. However, our data also support the idea that Baz is the weak link in maintaining epithelial integrity. In *pyd cno* mutants disruption of Baz localization is much more dramatic than effects on core AJ proteins. It is lost earlier in regions in which junctional remodeling is occurring and Baz localization became less uniform even in regions of the epidermis that remain intact.

Baz/Par3 has an interesting and complex relationship with myosin. During germband extension Baz and myosin have opposing planar polarization (reviewed in Kong *et al*., 2017), with activation of Rho kinase and Abelson tyrosine kinase (Abl) on AP borders working together to simultaneously activate myosin and downregulate cortical localization of AJ proteins and Baz (Simoes Sde *et al*., 2010; Tamada *et al*., 2012). The tension generated by myosin contractility reinforces these opposing polarities (Fernandez-Gonzalez *et al*., 2009; Fernandez-Gonzalez and Zallen, 2009; Levayer *et al*., 2011). However, during mesoderm apical constriction, tension generated by apical myosin contractility can also strengthen AJs (Weng and Wieschaus, 2016, 2017), suggesting the circuitry connecting adhesion and myosin-driven contractility is complex and context dependent. In one-cell *C. elegans* embryos Par3 and myosin have a similarly complex relationship. Par3 and contractile myosin foci both are found in the anterior domain but they are interspersed rather than co-localizing. Par proteins regulate myosin contractility (Cheeks *et al*., 2004; Munro *et al*., 2004), while cortical myosin flow and contractility regulating Par protein polarization and clustering (Dickinson *et al*., 2017; Rodriguez *et al*., 2017; Wang *et al*., 2017). Our data are especially intriguing given the known role of Baz as a regulator of apicomedial actomyosin contractility in the *Drosophila* amnioserosa, where it regulates pulsatile contractions and the degree of coupling to cell shape change, via effects on atypical protein kinase C (aPKC; David *et al*., 2010; Durney *et al*., 2018). Experiments in mammalian cells suggest Par3 may help regulate how cells balance apicomedial versus junctional contractility (Zihni *et al*., 2017). Par3 and ZO-1 are in proximity in mammalian cells (e.g. Van Itallie *et al*., 2013). How Cno and Pyd act to maintain Baz at AJs remains to be determined: is it via direct interactions or more indirect mechanisms? Together, these data support the idea that loss of cortical Baz in *cno* and *pyd cno* mutants may affect cortical contractility, but the complexity of the context-dependent roles of Baz/Par3 in different contexts leave questions to be answered in the future.

### Cno and Pyd: cooperative or parallel functions?

Another challenge for future work is defining the mechanisms by which Pyd/ZO-1 and Cno/Afadin act together and/or in parallel. Current data remain somewhat paradoxical. Both *Drosophila* and mammalian family members can directly bind one another. In mammals, ZO-1 is best known for its roles in tight junctions, while Afadin is an AJ protein, suggesting spatial separation—however, experiments using biotin-ligase proximity assessment in MDCK cells suggest at least some fraction of each protein are in close proximity (Van Itallie *et al*., 2013). In flies, both Cno and Pyd co-localize to AJs. Previous genetic assessments revealed genetic interactions, but these involved either partially functional alleles or knockdown (Yamamoto *et al*., 1997; Takahashi *et al*., 1998; Choi *et al*., 2016)—this rendered difficult assessment of whether an interaction involves two proteins acting in the same pathway or two proteins operating in separate but parallel pathways. Our current work uses embryos maternally and zygotically null for Pyd, avoiding this ambiguity.

The strong genetic enhancement of *pyd* null embryos by Cno knockdown suggests that the two proteins act, at least in part, in parallel. Another dataset supporting the idea that Cno and Pyd act in part in parallel rather than together comes from their respective localizations during germband extension—Pyd is planar polarized along with AJ proteins and Baz on DV boundaries (Suppl. Fig 5A,B) while Cno is enriched on AP boundaries with actin and myosin (Sawyer *et al*., 2011). Each could thus reinforce junctions at their respective locations, with combined loss further destabilizing adhesion. Alternately, they could have quite distinct functions: our work in MDCK cells suggests that ZO-1 family proteins can negatively regulate actomyosin assembly and contractility at AJs, by negatively regulating the Rok activator Shroom, while Afadin strengthens junction-cytoskeletal connections under tension (Choi *et al*., 2016). Intriguingly, Shroom plays a role in the reciprocal planar polarization of myosin and Baz, with its activity elevated at the borders where myosin is elevated and Baz and Pyd are reduced (Simoes Sde *et al*., 2014). Thus downregulation of Shroom and thus myosin contractility by Pyd is a possibility in *Drosophila*, though the relatively mild *pyd* and *shroom* single mutant phenotypes (Choi *et al*., 2011; Simoes Sde *et al*., 2014) suggest that in flies other proteins play overlapping and partially redundant roles. A different speculative possibility is that Afadin and ZO-1 proteins form part of a single large and multivalent protein complex, but recruit into this complex different, non-redundant partners, making loss of both more deleterious than loss of either one alone. Future work, including super-resolution imaging of Pyd and Cno, and identification and functional analysis of different binding partners may help resolve these mechanistic questions, and also help define what other proteins act in parallel or together with them—one intriguing candidate is the mechanosensitive junctional protein Jub/Ajuba (Razzell *et al*., 2018; Rauskolb *et al*., 2019) Defining interacting partners and parallel mechanisms for maintaining epithelial integrity is an important future direction.

## Materials and Methods

### Fly Stocks

Fly stocks are listed in Table 2. Wildtype was *yellow white*, *Histone-GFP or Histone-RFP*. Most experiments were performed at 25°C; we also carried out *cnoS-RNAi* at 27° to potentially increase severity of knockdown. Mutations are described at Flybase, https://flybase.org/. *shRNA* knockdown of *cno* was carried out by crossing double-copy *mat-tub-GAL4*, single-copy *mat-tub-Gal4*, *nos-Gal4*, or *MTD-Gal4* females, to males carrying UAS.*cnoValium20*shRNAi (control experiments for the *pyd* cno double mutants) or UASc*noValium22shRNA* constructs. The *pyd^B12^ cnoRNAiV20* chromosome (used to generate *pyd cno* double mutants) was generated through homologous recombination on chromosome 3 using the *pyd^B12^* deficiency allele (Choi et al., 2011) and the UAS.*cno Valium20* allele. *pyd^B12^ cnoRNAiV20* females were then crossed to male single-copy *mat-tub-Gal4; pyd^B12^/TM3* males.

**Table 2:** Fly Stocks

| Fly Stocks | Source |
| --- | --- |
| *UAS.cno RNAiV22* (stock #38194) | Bloomington Drosophila Stock Center (Bloomington, IL USA) |
| *UAS.cnoRNAiV20* (stock #33367) | Bloomington Drosophila Stock Center (Bloomington, IL USA) |
| *pyd<sup>B12</sup>/TM3* (stock # 51329) | Choi *et al.*, 2011 |
| *cno<sup>R2</sup>* (stock # 51328) | Sawyer *et al.*, 2009 |
| EcadGFP (stock #60584) | Bloomington Drosophila Stock Center (Bloomington, IL USA) |
| ZipGFP (stock #51564) | Bloomington Drosophila Stock Center (Bloomington, IL USA) |
| Nos-Gal4 (stock #32563) | Bloomington Drosophila Stock Center (Bloomington, IL USA) |
| MTD-Gal4 (nos-Gal4;nos-VP16-Gal4;otu-Gal4) (stock #31777) | Bloomington Drosophila Stock Center (Bloomington, IL USA) |
| Mat-tub-Gal4;Mat-tub-Gal4 (stock #80361) | Bloomington Drosophila Stock Center (Bloomington, IL USA) |
| Mat-tub-Gal4 (stock #7062) | Bloomington Drosophila Stock Center (Bloomington, IL USA) |

### Cuticle preparation

Cuticle preparation was performed according to Wieschaus and Nüsslein-Volhard, (1986) with the following modifications. Embryos were collected and allowed to develop for 48 hrs at 25°C. Unhatched embryos were washed in 0.1% Triton X-100 and dechorionated in 50% bleach for 5 minutes in a glass depression slide. After washing once in 0.1% Triton X-100, embryos were mounted in 1:1 diluted Hoyer’s medium:lactic acid and incubated at 60°C overnight.

### Immunofluorescence

Antibodies and dilutions used are listed in Table 3. All embryos were dechorionated in 50% bleach for 5 minutes on nutator. Most antibody staining was done on “heat fixed” embryos, which were fixed in boiling Triton salt solution (0.03% Triton X-100, 68 mM NaCl, 8 mM EGTA) for 10 seconds followed by fast cooling on ice and devitellinized by vigorous shaking in 1:1 heptane:methanol. Embryos were stored in 95% methanol/5% 0.5 M EGTA for at least 48 hours at −20°C prior to staining. Embryos were washed three times in blocking solution (0.01% Triton X-100 in PBS; 0.01% PBS-T) with 1% normal goat serum (NGS), followed by blocking during nutation for 1 hour. For visualization of F-actin with Phalloidin, embryos were Formaldehyde fixed in 1:1 4% formaldehyde in phosphate-buffered saline (PBS):heptane for 20 minutes or 1:1 8% formaldehyde in phosphate-buffered saline (PBS):heptane for 30 minutes, while rocking. Embryos were devitellinized by vigorous shaking in 1:1 heptane:methanol,, except when visualizing F-actin using Phalloidin, when they were devitellinized using 1:1 heptane:95% ethanol by manual removal of the vitelline membrane with a scalpel. Primary and secondary antibody staining were each carried out at 4°C overnight with nutation in 1% NGS in PBS-T. After primary and secondary antibody staining the embryos were washed three times for 5 minutes with 0.01% Triton X-100 in PBS (PBS-T).

**Table 3:** Antibodies

| **Immunofluorescence** |  |  |
| --- | --- | --- |
| Antibodies | Dilution | Source |
| **Primary Antibodies** |  |  |
| Anti-Canoe (rabbit) | 1:2000 | J. Sawyer and N. Harris (UNC, Chapel Hill, NC, USA) |
| Anti-DE-cadherin (rat) | 1:50 | Developmental Studies Hybridoma Bank (DSHB),( University of Iowa, Iowa City, IA, USA) |
| Anti-Armadillo (mouse) | 1:100 | DSHB |
| Anti-Neurotactin (mouse) | 1:100 | DSHB |
| Anti-Enabled (mouse) | 1:500 | DSHB |
| Anti-Polychaetoid (mouse) | 1:250 | Choi *et al.*, 2011 |
| Anti-Bazooka (rabbit) | 1:2000 | Choi *et al.*, 2013 |
| Anti-Myosin II Heavy Chain (rabbit) | 1:500 | Chougule *et al.*, 2016 |
| **Secondary Antibodies** |  |  |
| AlexaFluor 405,488,568, and 647 | 1:500 | Life Technologies |
| Phalloidin TRITC and FITC | 1:500 | Thermo Fisher Scientific |
| **Immunoblotting** |  |  |
| **Primary Antibodies** |  |  |
| Anti-Armadillo (mouse) | 1:500 | DSHB |
| Anti-Canoe (rabbit) | 1:1000 | J. Sawyer and N. Harris (UNC, Chapel Hill, NC, USA) |
| Anti-Polychaetoid (mouse) | 1:1000 | Choi *et al.*, 2011 |
| Anti-αTubulin (mouse) | 1:5000 | Sigma Life Science, T6199 |
| **Secondary Antibodies** |  |  |
| IRDye 680RD (anti-rabbit) | 1:10000 | LI-COR Biosciences |
| IRDye 800CW (anti-mouse) | 1:10000 | LI-COR Biosciences |

### Image acquisition and manipulation

Fixed embryos were mounted in Aqua-Poly/Mount (Polysciences) and imaged on a confocal laser-scanning microscope (LSM 710 or LSM880, 40×/NA 1.3 Plan-Apochromat oil objective, Carl Zeiss). ZEN 2009 software (Carl Zeiss) was used to process images and render *z*-stacks in 3D. Super resolution embryos were imaged on Nikon N-SIM, using SR APO TIRF 100X/1.49 NA oil objective. NIS-Elements AR Version 4.51 software 2016 (Nikon) was used to process SIM images and render z-stacks in 3D. Photoshop CS6 (Adobe) was used to adjust input levels so that the signal spanned the entire output grayscale and to adjust brightness and contrast. Image processing for SIM imaging is described in Suppl. Fig. 2.

### Immunoblotting

Antibodies and dilutions used are listed in Table 3. Expression levels of Arm protein and knockdown efficiency of Cno and Pyd were measured by immunoblotting embryos of 1-4 and 12-15 hours old. Embryo lysates were performed according to Bonello *et al.*, 2018 with the following modification. The lysis buffer was 1% NP-40, 0.5% Na deoxycholate, 0.1% SDS, 50 mM Tris pH 8.0, 300 mM NaCl, 1.0 mM DTT, Halt protease and phosphatase inhibitor cocktail and 1 mM EDTA. Lysates were resolved using a 7 or 8% SDS-PAGE and transferred to nitrocellulose membrane. The membranes were incubated with the primary antibody for 2 hours at room temperature or overnight at 4°C (see Table# for antibody concentrations). The membranes were incubated with secondary antibody for 45 minutes at room temperature (see Table# for antibody concentrations). The signal levels were detected using Odyssey CLx infrared system (LI-COR Biosciences). Band densitometry was calculated using LI-COR Image Studio.

## Acknowledgements

We are grateful to Zachary Blom and Wangsun Choi for initiating the project, to Teresa Bonello for carrying out myosin staining, to Melissa Greene, Clara Williams and Halle Ronk for technical assistance, to Ulli Tepass, Sergio Simoes, Paul Maddox, Teresa Bonello, Amy Gladfelter, Kevin Slep, and Kathryn Reissner for helpful discussions, to the Bloomington Drosophila Stock Center, Tony Perdue for advice on SIM and Confocal imaging, to the Bloomington Drosophila Stock Center, Jeffrey Thomas at Texas Tech University Health Sciences Center for Myosin II Heavy Chain primary antibody, Michelle Itano of the Neuroscience Center Microscopy Core and Pablo Ariel of the Microscopy Services Laboratory for advice on 3D imaging, and to members of the Peifer lab for helpful discussions and reading the manuscript. The work was supported by NIH R35 GM118096 to M.P. L.A.M. was supported by NIH K12 GM000678, K.Z.P-V. by NIH R25 GM055336 and T32 GM007092, and M.T.S. by a Diversity Supplement on NIH R35 GM118096. M.T.S. was a participant in the UNC PREP Program, which is funded by NIH R25 GM089569.

