## Supplemental Figures for "The Drosophila Afadin and ZO-1 homologs Canoe and Polychaetoid act in parallel to maintain epithelial integrity when challenged by adherens junction remodeling"

Suppl. Fig 1. Efficiency of Cno knockdown after *cnoS-RNAi* and in *cno pyd* double mutants, and effects on Pyd and Arm levels. A-F. Immunoblots revealing the levels of Cno, Pyd, and Arm and quantification of Cno levels. Wildtype embryos serve as the baseline for quantification, and thus effect of knockdown or mutations on levels of Cno, Pyd and Arm are normalized to wildtype. Antibodies used to probe the membranes are indicated at the right and molecular weight standards are indicated on the left. The age of the embryos analyzed is located below the Western blot. $\alpha$-tubulin served as a loading control. A. Levels of Cno, Pyd, and Arm in wildtype versus those from the *cnoS-RNAi* cross. Note there are multiple isoforms of Pyd which vary in relative levels over developmental time (Choi *et al.,* 2011). B,C. Quantification of Cno protein levels. (B) In early embryos (1-4 h) Cno is highly reduced compared to wild type—quantification suggested it was reduced 1.3±0.6% but this may simply reflect background. C. During late embryogenesis (12-15 h), Cno levels are still highly downregulated compared to wild type (4.85%). D. Levels of Cno, Pyd, and Arm in wildtype, from the *pyd, cno* double mutant cross, and from the *cnoshRNAv20* cross used as our *cno* control. E,F. Quantification of Cno protein levels. E. In early embryos (1-4 h), Cno is highly reduced compared to wildtype in both the *cno* control cross (3.6±1.0%) and the *pyd cno* cross (2.7±0.00%). F. During late embryogenesis (12-15 h), Cno levels remain downregulated compared to wildtype in both the *cno* control cross (8.1±0.65%) and the *pyd cno* cross (6.5±1.0%). G,H. Stage 9 wildtype and *cnoS-RNAi* embryos stained and imaged together. Cno staining is dramatically reduced. I,J. Stage 10 wildtype and *pyd cno* embryos stained and imaged together. Cno staining is dramatically reduced, and Pyd staining is absent (this was one of the 50% of the embryos that show no Pyd staining and thus are presumptive *pydMZ* mutants.) Scale bars=30µm.

Suppl. Fig. 2. Methodology used to process structured illumination (SIM) super-resolution images. A. Wildtype Confocal image of a mid-stage 13 (mid dorsal closure) embryo. Ena puncta localize to tricellular junctions with Actin highly enriched at the leading edge. B. Wildtype SIM image of mid-stage 13 (mid stage dorsal closure). SIM allows for higher resolution of Ena and Actin. C. Look Up Table (LUT) for image in Panel B, which displays intensity histogram. Most pixels in the embryo show essentially background accumulation, and thus have little relevant information despite apparent differences in intensity. In processing SIM images, we examine the brightest subset of the pixels, which include those documenting cortical Ena, actin or AJ protein localization. Dotted lines here illustrate the boundaries of the display range used to generate the image shown in Panel B. D. 3-D N-SIM utilized 5 different phases and 3 different orientations (angles) of the interference pattern. This results in a 3 X 5 image. E. Look Up Table (LUT) for the image in Panel D. F. NIS Computational reconstruction of the image in panel D. G. Look Up Table (LUT) for image in Panel F, with dotted lines indicating boundaries used by the reconstruction software. H. Final SIM image like those presented in Figure 5. I. Look Up Table (LUT) for the image in Panel H, with dotted lines indicating boundaries used. All scale bars=20µm

Suppl. Fig. 3. Crosses used to generate *cnoS-RNAi, pyd cno*, and *cno* control cross embryos. *cnoS-RNAi* provided our strongest loss of Cno. The cross used as a control for the *pyd cno* double mutants was chosen to provide the same degree of Cno knockdown as in the *pyd cno* double mutant cross. Offspring genotypes leave out most GAL4 drivers as all drive expression exclusively maternally and thus their presence or absence in the embryo is irrelevant for phenotypic strength.

Suppl. Fig. 4. *cnoS-RNAi* mimics effects of *cno* maternal/zygotic mutants on early cytoskeletal and junctional planar polarity. A,B,G,H. Stage 7 embryos just after the start of germband extension. A,G. In wildtype Baz becomes mildly planar-polarized, with enrichment at dorsal/ventral cell borders (blue arrows) relative anterior/posterior cell borders (yellow arrows). B,H. In *cnoS-RNAi* embryos, Baz planar polarity is enhanced and Baz localization is more punctate. C-F,I,J. Stage 8 embryos. C,I. In wildtype, myosin becomes planar polarized in an opposite fashion to Baz, accumulating at higher levels on anterior/posterior borders (C,I inset, arrows). D,I. In *cnoS-RNAi* embryos these planar-polarized myosin cables separate from the AJ on anterior-posterior borders (blue arrows) and at multicellular junctions at the center of rosettes (yellow arrows). E,F,J. During stage 8 mitotic domains began firing, with mitotic domain 11 most prominent in the thorax and abdomen. Cortical Arm is reduced in intensity on cells that have rounded up to divide (arrows). This occurs at the correct time in *cnoS-RNAi* embryos. Scale bars =20µm.

Suppl. Fig. 5. Pyd planar polarity tracks with that of AJ proteins and Cno loss does not dramatically alter Pyd cortical localization. A,B. Stage 8 embryos, when planar polarity is most apparent. A. In wildtype planar polarity of AJ proteins is quite subtle, with that of Baz most apparent, with elevated levels on DV (blue arrows) relative to AP boundaries (yellow arrows). B. Planar polarization of Arm, Baz and Pyd is accentuated after *cnoS-RNAi*, with the effects on Baz being the most dramatic, with strong reduction on AP (yellow arrows) relative to DV borders (blue arrows). C,D. Stage 9. Pyd localization continues to parallel that of AJ proteins in both wildtype and *cnoS-RNAi*, at a stage when Baz localization is more severely altered. Scale bars=30µm.

Suppl. Fig 6. The most strongly affected embryos from the *cno* control cross phenocopy those after *cnoS-RNAi*. Progeny of *matGAL4/+; cnov20shRNA /+* females x *cnov20shRNA /+* males. A-B”. Mid Stage 9. Dorsal ectodermal cells (magenta brackets) have competed their first mitotic divisions, resumed columnar architecture and are hyperconstricted, similar to what we observed after *cnoS-RNAi* (Fig. 6D). Most cells in the dorsal neuroectoderm remain rounded up after division (yellow brackets), with reduced cortical Baz. Ventral neurectoderm (cyan brackets) remains intact prior to divisions. C-D”. Late Stage 9. Cell in the M mitotic domain are dividing (arrows) across the ventral ectoderm (cyan brackets). Cells in the neuroectoderm have reduced and fragmented cortical Baz (blue brackets), similar to *cnoS-RNAi* (Fig. 6J). E-F” Stage 10. Many cells in the ventral neurectoderm, especially along the midline (blue brackets), have lost epithelial integrity and have highly reduced cortical Baz, similar to *cnoS-RNAi* (Fig. 6L). Scale bars= 20 µm.

Suppl. Fig. 7. *pyd cno* mutants share early defects with *cnoS-RNAi.* A. Cellularizing embryos, maximum intensity projections (MIPs) of cross sections. Top. In wildtype Baz localizes to nascent AJs while Arm is in both apical spot AJs and basal junctions. Middle and bottom. In both *cnoS-RNAi* and *pyd cno* embryos, Baz and Arm enrichment in nascent AJs is lost, while basal junctions remain. Baz localizes to nascent AJs while Arm is in both apical spot AJs and basal junctions. B,C. Stage 7 embryos. Mesoderm fails to full invaginate with myosin forming apical balls in mesodermal cells, as we previously observed in *cnoMZ* mutants. C. Baz and Arm become hyper-planar polarized. Large grooves form at sites of the dorsal folds (yellow arrows). Ectodermal cells align in rows (magenta arrows) with gaps appearing along AP cell borders (blue arrows). D,E. Stage 7. Cortical myosin accumulation increases on AP cell borders and it separates from the cell junctions (E, arrows). Scale bars=10µm.

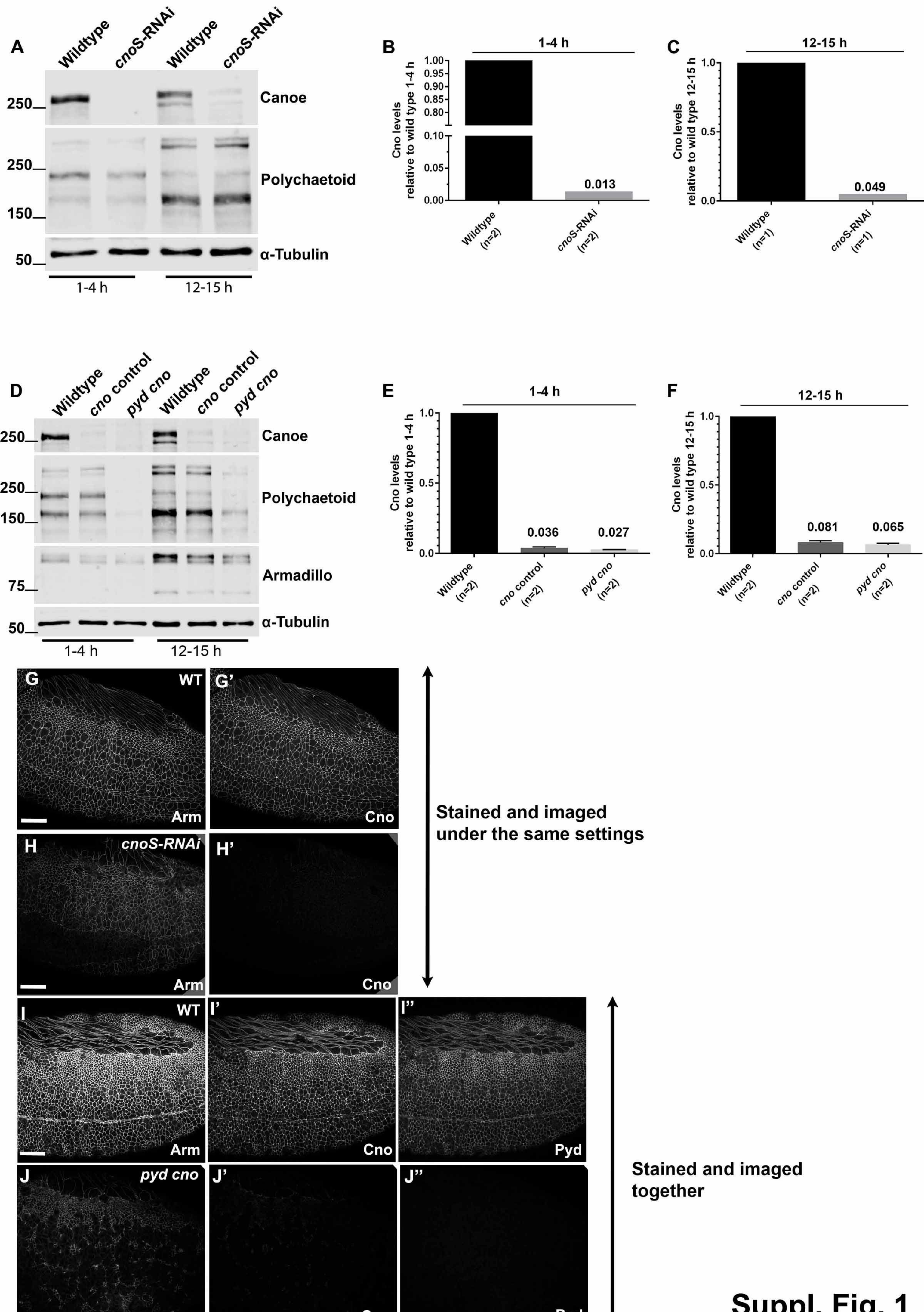
A
Wildtype
cnoS-RNAi
Wildtype
cnoS-RNAi
250
Canoe
250
150
Polychaetoid
50
α-Tubulin
1-4 h
12-15 h
B
1-4 h
Cno levels relative to wild type 1-4 h
0.013
Wildtype (n=2)
cnoS-RNAi (n=2)
C
12-15 h
Cno levels relative to wild type 12-15 h
0.049
Wildtype (n=1)
cnoS-RNAi (n=1)
D
Wildtype
cno control
pyd cno
Wildtype
cno control
pyd cno
Canoe
Polychaetoid
Armadillo
α-Tubulin
75
1-4 h
12-15 h
E
1-4 h
Cno levels relative to wild type 1-4 h
0.036
0.027
Wildtype (n=2)
cno control (n=2)
pyd cno (n=2)
F
12-15 h
Cno levels relative to wild type 12-15 h
0.081
0.065
Wildtype (n=2)
cno control (n=2)
pyd cno (n=2)
G
WT
G'
Arm
Cno
H
cnoS-RNAi
H'
Arm
Cno
Stained and imaged under the same settings
I
WT
I'
I''
Arm
Cno
Pyd
J
pyd cno
J'
J''
Arm
Cno
Pyd
Stained and imaged together

Suppl. Fig. 1

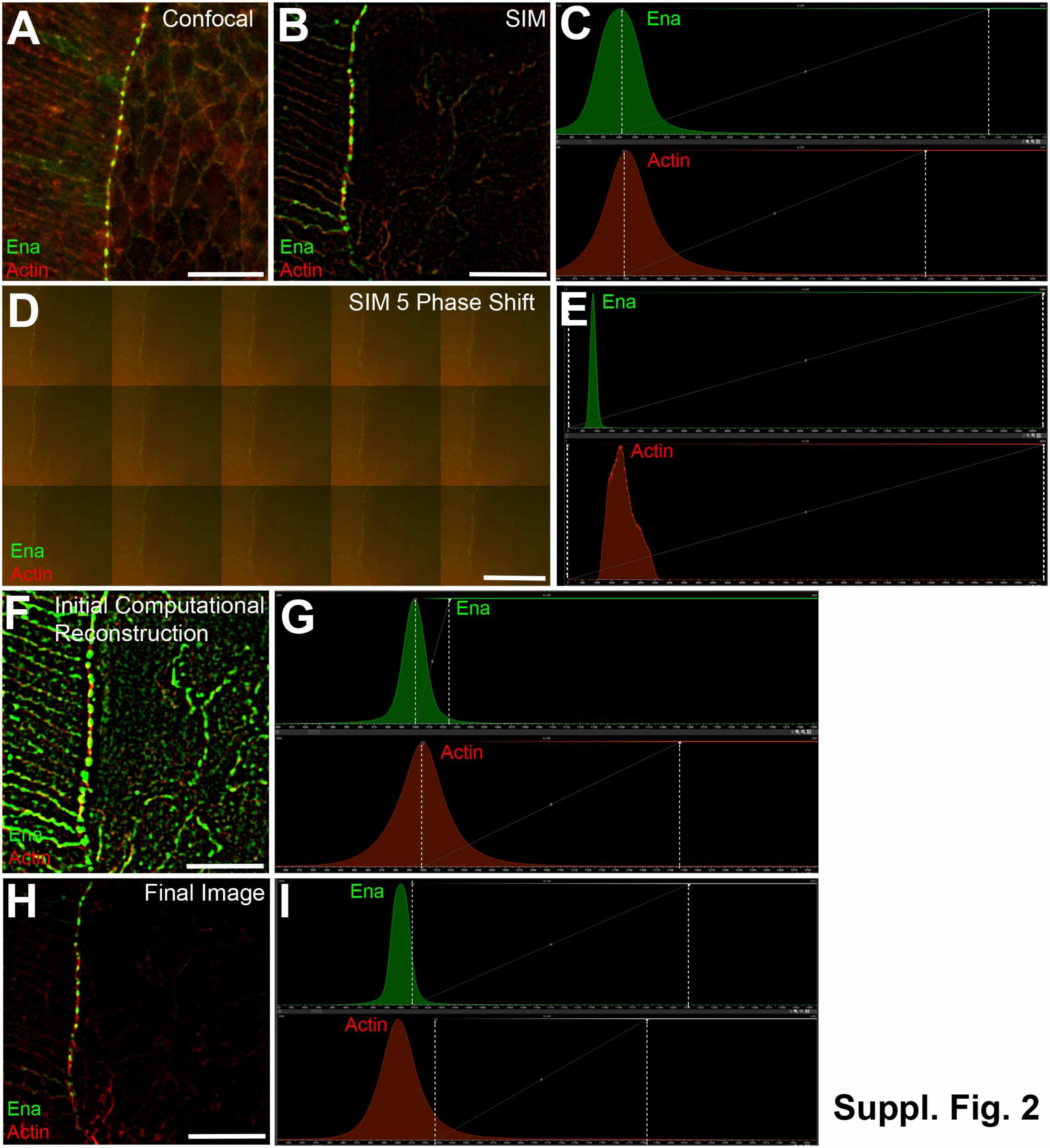

A
Confocal
Ena
Actin
B
SIM
Ena
Actin
C
Ena
Actin
D
SIM 5 Phase Shift
Ena
Actin
E
Ena
Actin
F
Initial Computational Reconstruction
Ena
Actin
G
Ena
Actin
H
Final Image
Ena
Actin
I
Ena
Actin


Suppl. Fig. 2

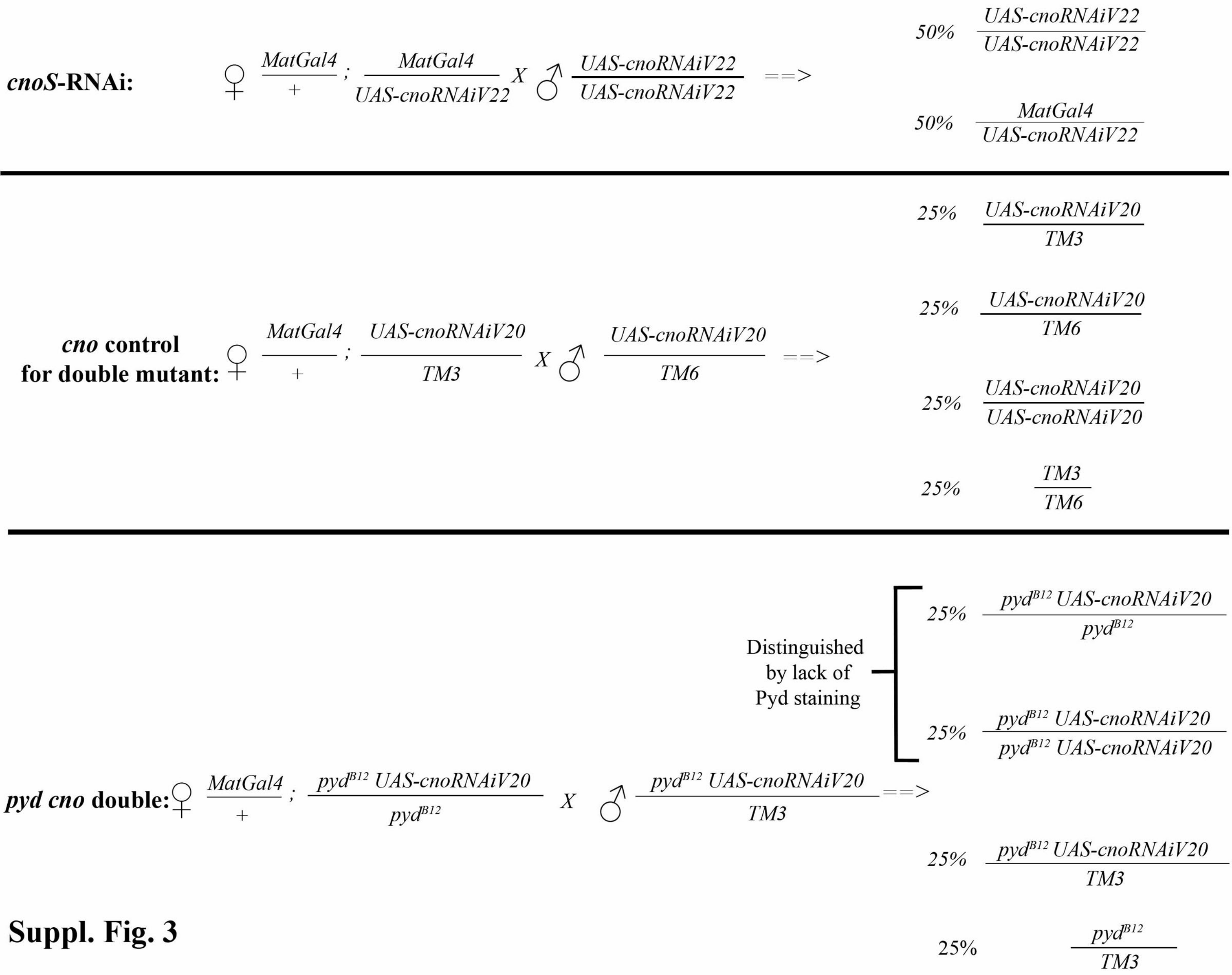

cnoS-RNAi:
♀ MatGal4 / + ; MatGal4 / UAS-cnoRNAiV22 X ♂ UAS-cnoRNAiV22 / UAS-cnoRNAiV22 ==>
50% UAS-cnoRNAiV22 / UAS-cnoRNAiV22
50% MatGal4 / UAS-cnoRNAiV22
cno control for double mutant:
♀ MatGal4 / + ; UAS-cnoRNAiV20 / TM3 X ♂ UAS-cnoRNAiV20 / TM6 ==>
25% UAS-cnoRNAiV20 / TM3
25% UAS-cnoRNAiV20 / TM6
25% UAS-cnoRNAiV20 / UAS-cnoRNAiV20
25% TM3 / TM6
pyd cno double:
♀ MatGal4 / + ; pydB12 UAS-cnoRNAiV20 / pydB12 X ♂ pydB12 UAS-cnoRNAiV20 / TM3 ==>
Distinguished by lack of Pyd staining
25% pydB12 UAS-cnoRNAiV20 / pydB12
25% pydB12 UAS-cnoRNAiV20 / pydB12 UAS-cnoRNAiV20
25% pydB12 UAS-cnoRNAiV20 / TM3
25% pydB12 / TM3


**Suppl. Fig. 3**

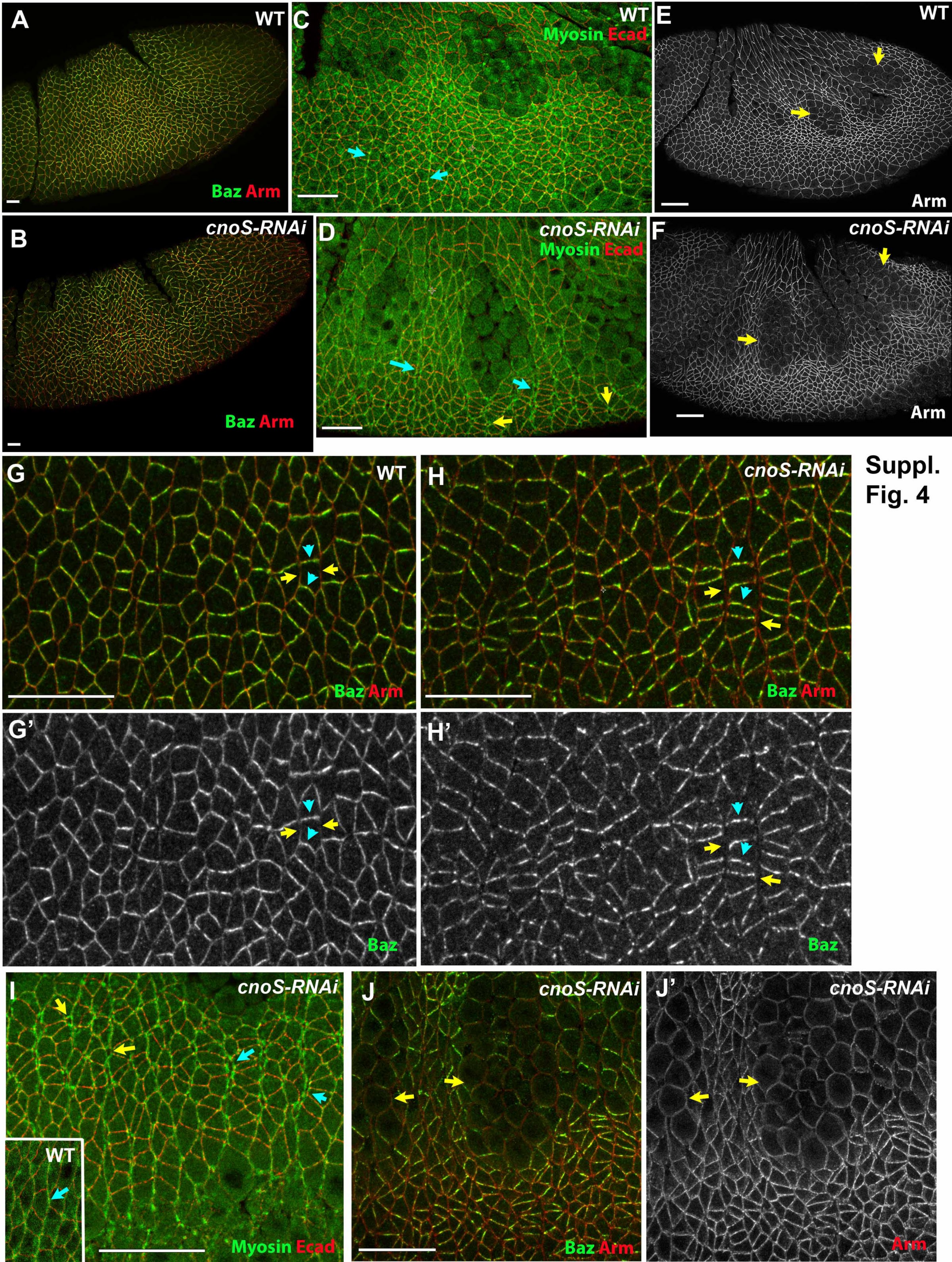
A
WT
Baz Arm
B
cnoS-RNAi
Baz Arm
C
WT
Myosin Ecad
D
cnoS-RNAi
Myosin Ecad
E
WT
Arm
F
cnoS-RNAi
Arm
G
WT
Baz Arm
H
cnoS-RNAi
Baz Arm
G'
Baz
H'
Baz
I
cnoS-RNAi
WT
Myosin Ecad
J
cnoS-RNAi
Baz Arm
J'
cnoS-RNAi
Arm
Suppl.
Fig. 4

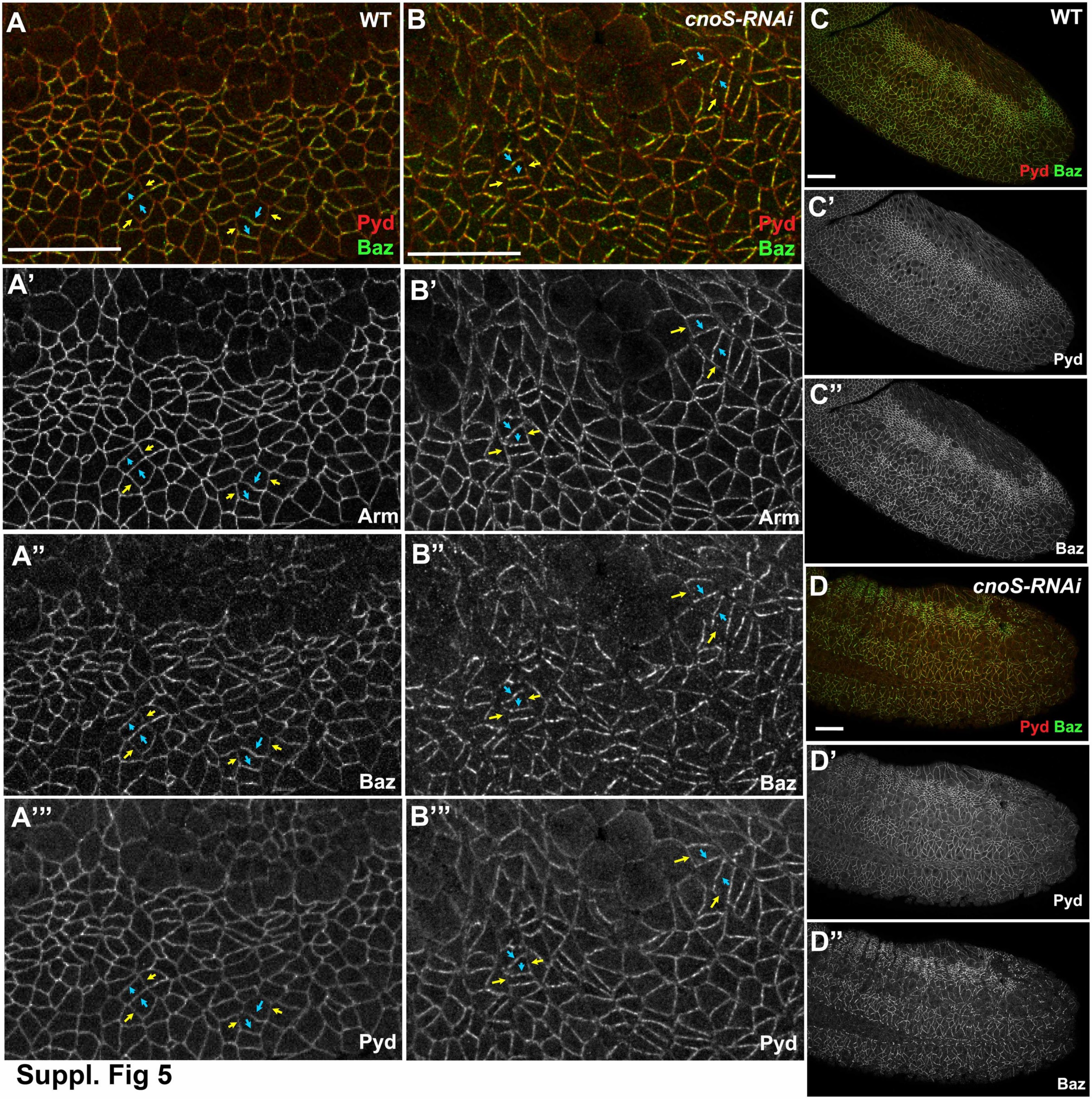
A
WT
Pyd
Baz
B
cnoS-RNAi
Pyd
Baz
C
WT
Pyd Baz
C'
Pyd
C''
Baz
A'
Arm
B'
Arm
A''
Baz
B''
Baz
D
cnoS-RNAi
Pyd Baz
D'
Pyd
D''
Baz
A'''
Pyd
B'''
Pyd

Suppl. Fig 5

**Suppl. Fig 6**

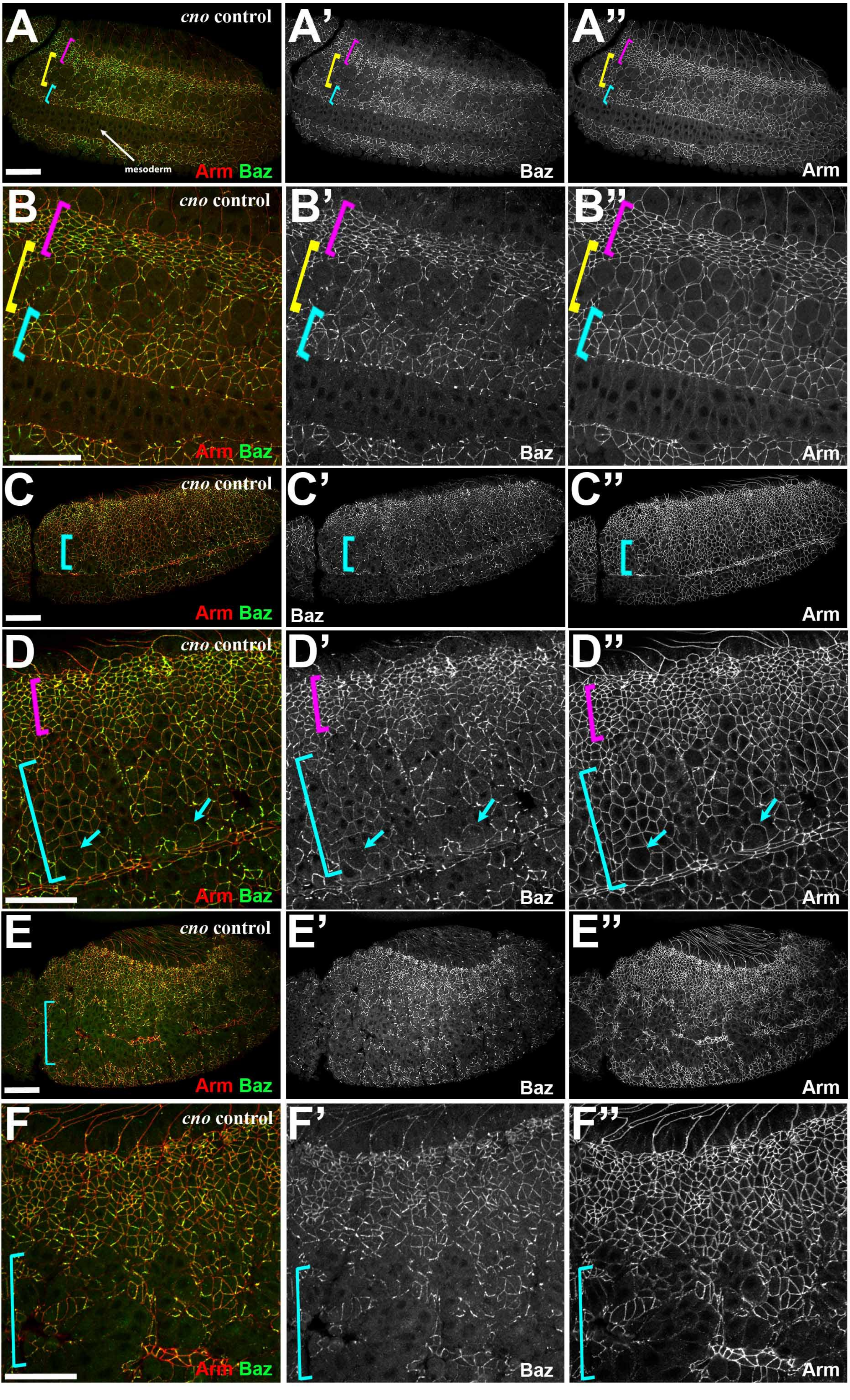

A
*cno* control
mesoderm
Arm Baz
A'
Baz
A''
Arm
B
*cno* control
Arm Baz
B'
Baz
B''
Arm
C
*cno* control
Arm Baz
C'
Baz
C''
Arm
D
*cno* control
Arm Baz
D'
Baz
D''
Arm
E
*cno* control
Arm Baz
E'
Baz
E''
Arm
F
*cno* control
Arm Baz
F'
Baz
F''
Arm

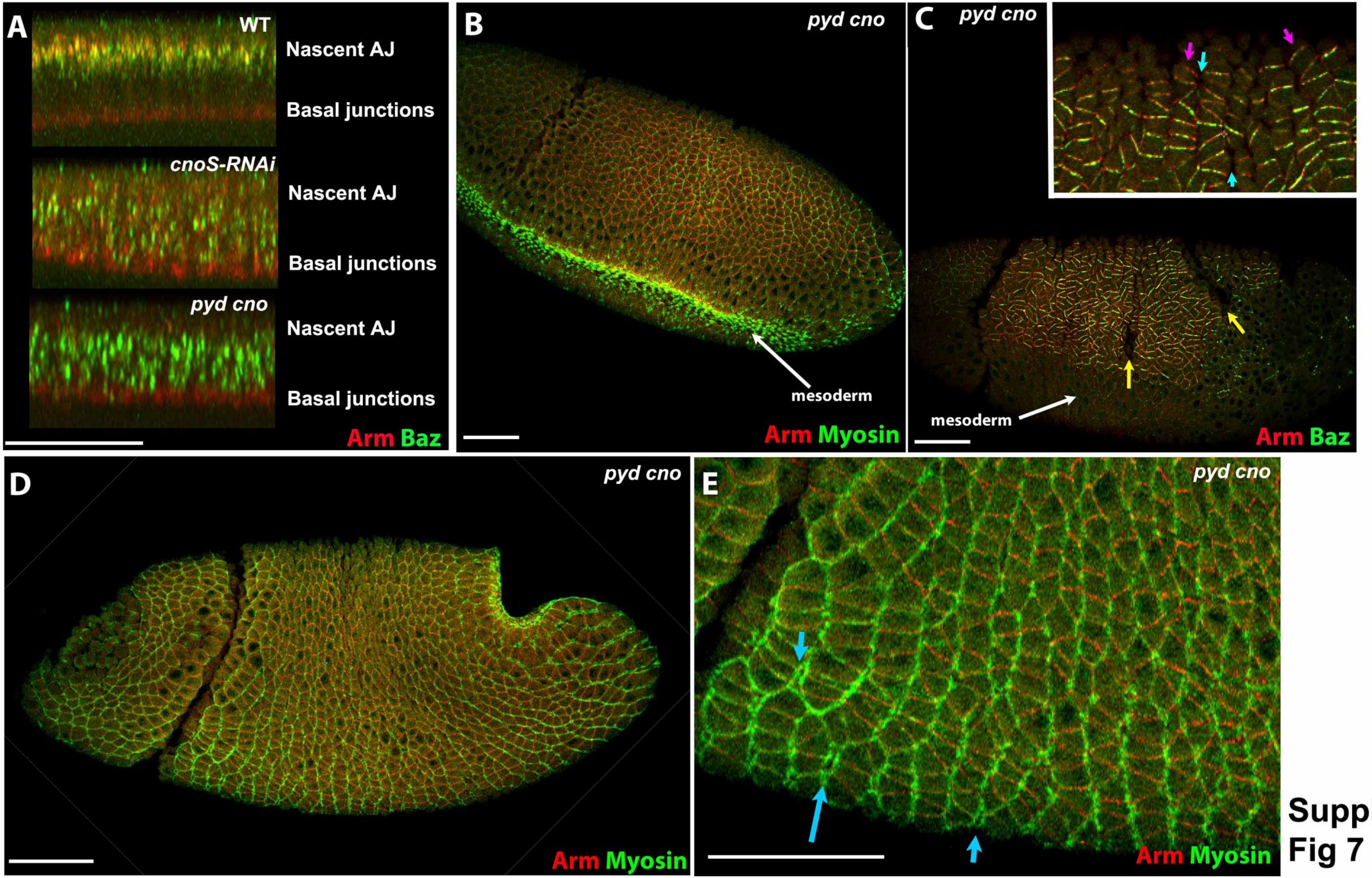
A
WT
Nascent AJ
Basal junctions
cnoS-RNAi
Nascent AJ
Basal junctions
pyd cno
Nascent AJ
Basal junctions
Arm Baz
B
pyd cno
mesoderm
Arm Myosin
C
pyd cno
mesoderm
Arm Baz
D
pyd cno
Arm Myosin
E
pyd cno
Arm Myosin

Suppl. Fig 7
